# Cell-type-specific roles of inhibitory interneurons in the rehabilitation of auditory cortex after peripheral damage

**DOI:** 10.1101/2022.09.15.508128

**Authors:** Manoj Kumar, Gregory Handy, Stylianos Kouvaros, Lovisa Ljungqvist Brinson, Brandon Bizup, Brent Doiron, Thanos Tzounopoulos

## Abstract

Peripheral sensory organ damage leads to compensatory cortical plasticity that supports a remarkable recovery of perceptual capabilities. A major knowledge gap is the lack of precise mechanisms that explain how this plasticity is implemented and distributed over a diverse collection of excitatory and inhibitory cortical neurons. Here, we explored these mechanisms in mouse A1. After peripheral damage, we found recovered sound-evoked activity of excitatory principal neurons (PNs) and parvalbumin (PVs) interneurons (INs), reduced activity in somatostatin-INs (SOMs), and recovered activity in vasoactive intestinal peptide-INs (VIPs). Given the sequentially organized cortical network where VIPs inhibit INs, SOMs inhibit PVs and PNs, and PVs inhibit PNs, our results suggest that PVs contribute to PN stability, SOMs allow for increased PN and PV activity, and VIPs enable the PN and PV recovery by inhibiting SOMs. These results highlight a strategic, cooperative, and cell-type-specific plasticity program that restores cortical sound processing after peripheral damage.

## Introduction

In all sensory systems, damage to peripheral organs leads to compensatory cortical reorganization and increased cortical sensitivity to the non-damaged (spared) sensory input^1-9^. This plasticity is crucial for survival, for it supports a remarkable recovery of perceptual capabilities^10,11^. Despite the great importance of this plasticity, the underlying system, circuit, and cellular mechanisms remain poorly understood. The establishment of these mechanisms will reveal major concepts in cellular and functional cortical rehabilitation after peripheral damage. Moreover, it holds the promise to highlight novel strategies for enhancing perceptual recovery and mitigating brain disorders associated with sensory deficits and subsequent pathogenic cortical plasticity, such as schizophrenia, tinnitus, phantom limb pain, and neuropathic pain^12-16^.

In the auditory system, while the auditory nerve input to the brainstem is significantly reduced after cochlear damage, the cortical sound-evoked activity is maintained or even enhanced^16-18^, due to increased cortical gain, the sensitivity of neuronal responses against sound levels. As such, this plasticity contributes to the recovery of perceptual sound-detection thresholds after cochlear damage^10,11,19-21^. The increased cortical gain is associated with reduced inhibitory (GABAergic) cortical activity, increased spontaneous firing, and reorganization of frequency tuning towards less damaged regions of the cochlea^6,10,18,22-26^. Moreover, a steep drop in PV-mediated inhibition to principal neurons (PNs) is a predictor of auditory cortical response rehabilitation after cochlear nerve damage^22^. Although the role of general or PV-centric reduced inhibition is well documented^11,22,24,26-29^, it does not provide the precise cellular and circuit mechanisms that mediate cortical rehabilitation.

The recent use of cell-type-specific labeling and optogenetic manipulations^30^, combined with the genetic and physiological dissection of cortical interneurons (INs)^31,32^, have established a new picture of our understanding of cortical circuits. The canonical cortical circuit includes (at a minimum) vasoactive intestinal-peptide (VIP), somatostatin (SOM), and parvalbumin (PV) expressing IN sub-classes, all with distinct and sequentially organized synaptic connections among themselves and PNs^33,34^. This circuit design begs for a more precise mechanistic understanding of how specific cortical gain modulations associated with an overall and non-specific decrease in inhibition, such as increased cortical gain after peripheral trauma, are implemented and distributed over these distinct IN sub-classes. Namely, cortical inhibition is crucial for suppressing neuronal activity^35-38^, firing rate gain modulation^39-44^, and spike timing control^45,46^, as well as for correlated neuronal^47,48^ and population activity^49,50^. Cortical inhibition is also essential for the prevention of runaway cortical activity that would otherwise lead to pathologic activity^37,51,52^. As such, this complex role of inhibition is expected to pose constraints on how reduced inhibition can safely modulate cortical gain^53^, for a global and non-specific inhibitory reduction could lead to instability and pathology, such as epileptic-like activity^52^.

To study the precise mechanisms of inhibition in cortical plasticity after peripheral damage, we used a mouse model of noise-induced cochlear damage. We employed electrophysiological and immunohistochemical assays to assess peripheral damage, behavioral assays to assess perceptual hearing thresholds, longitudinal *in vivo* two-photon calcium imaging to assess the activity of different cortical neuronal types, *ex vivo* electrophysiology assays to assess cellular excitability, and computational models to shape our hypotheses and predictions. Our results demonstrate that the recovery of cortical sensory processing after peripheral damage is supported by a remarkable cell-type-specific contribution and cooperativity among multiple types of cortical INs.

## Results

To cause peripheral damage in mice, we used a noise-induced hearing loss (NIHL) paradigm. Mice were bilaterally exposed to an octave band (8-16 kHz) noise at 100 dB SPL for 2 hours (**Fig. 1a, b)**. To assess the consequences of this noise exposure on peripheral structures, we measured and quantified the auditory brainstem response (ABR) before and one, three, and ten days after noise exposure. ABR represents the sound-evoked action potentials generated by the synchronized activity of various nuclei of the auditory pathway from the auditory nerve to the brainstem, where ABR wave 1 represents the sound-evoked synchronized activity of the auditory nerve (AN) type-I spiral ganglion neurons (SGNs) (**Fig. 1c**). We found that noise exposure increased the ABR threshold, the sound-level which elicited a significant wave 1 amplitude (**Fig. 1c-e**) and reduced the gain of that AN sound-evoked activity, the slope of ABR wave I amplitude against sound level, (**Fig. 1f, g**), suggesting reduced sound information relayed from the cochlea to the AN. Moreover, we found that noise exposure increased the distortion product otoacoustic emissions (DPOAE) threshold (**Fig. 1h**), suggesting dysfunction of the cochlear outer hair cells (OHCs) sound amplification function. ABR threshold and gain represent the combined functionality of inner hair cells (IHCs), OHCs, type-I SGNs, and synapses between the IHCs and type-I SGN dendrites called ribbon synapses^54^. To identify the anatomical markers of reduced AN gain and elevated ABR and DPOAE thresholds after noise exposure, we performed immunohistochemical analysis across the tonotopic axis of the cochlea to quantify the survival of IHCs, OHCs, and the number of ribbon synapses between the IHCs and type-I SGN dendrites (**Fig. 1i and supplement Fig. 1**). We found that noise exposure significantly reduced the number of ribbon synapses per inner hair cell in the high-frequency region (16-32 kHz) of the cochlea (**Fig. 1i, j**), without affecting the survival of either IHCs or OHCs (**supplement Fig. 1**). We did not observe any changes in sham-exposed mice, which underwent identical procedures but without the presentation of sound (**Fig. 1 and supplement Fig. 1**). Together, our noise trauma protocol, by reducing the AN gain and increasing peripheral hearing thresholds, reduces the amount and transfer of peripheral auditory input to the brain. We will use this protocol to assess perceptual recovery and the cellular mechanisms of cortical rehabilitation after peripheral damage.

**Figure 1.**
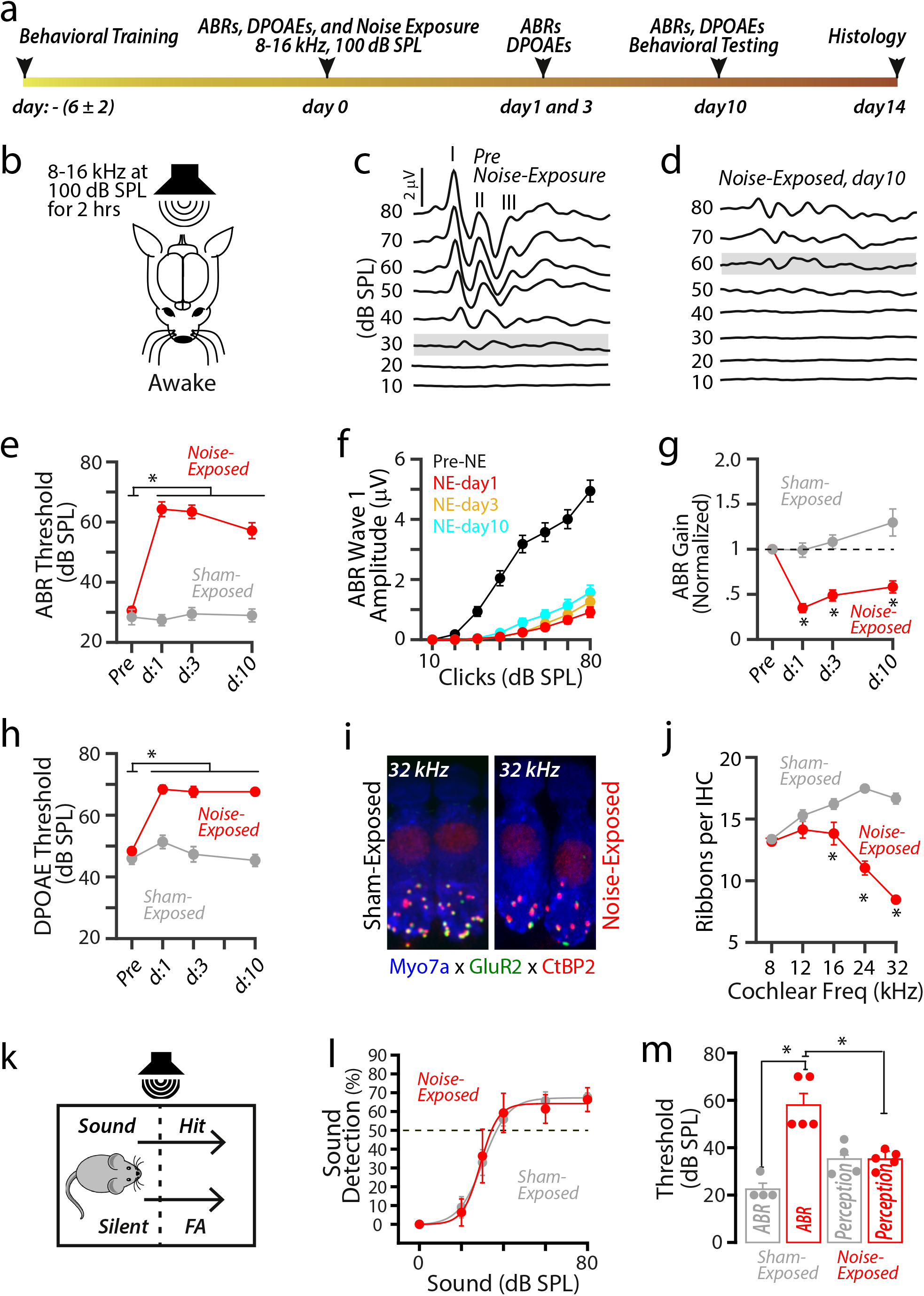
Perceptual sound-detection threshold recovers despite elevated ABR threshold after NIHL. **(a)** Timetable of experimental design. **(b)** Noise-exposure paradigm. **(c-d)** Representative ABR traces to clicks before and 10 days after noise exposure. **(e)** Average ABR thresholds from noise-exposed (n = 35) and sham-exposed (n = 19) mice, before and 1, 3, and 10 days after exposure. (Noise vs. sham: 2-way ANOVA; exposure x time interaction, F = 21.7, p = 2.8 × 10^−12^; effect of exposure, F = 221.3, p = 1.4 × 10^−34^; *, p < .05, compared to pre-noise-exposed, Holm-Bonferroni’s post hoc). **(f)** Average ABR wave 1 amplitude to clicks before and after noise exposure. (1-way repeated measure ANOVA; F = 227.7, p = 4.9 × 10^−38^; *, p < .05, compared to pre-noise-exposed, Holm-Bonferroni’s post hoc). **(g)** Average ABR gain from noise-exposed and sham-exposed mice, before and after exposure. (Noise vs. sham: 2-way ANOVA; exposure x time interaction, F = 11.3, p = 6.3 × 10^−7^; effect of exposure, F = 99.8, p = 1.8 × 10^−19^; *, p < .05, compared to pre-noise-exposed, Holm-Bonferroni’s post hoc). **(h)** Average DPOAE thresholds from noise-exposed (n = 5) and sham-exposed mice (n = 4), before and after exposure. (Noise vs. sham: 2-way ANOVA; exposure x time interaction, F = 10.9, p = 1.4 × 10^−6^; effect of exposure, F = 145.6, p = 1.9 × 10^−23^; *, p < .05, compared to pre-noise-exposed, Holm-Bonferroni’s post hoc). **(i)** Cochlear histology images of a 32 kHz frequency region from noise-exposed mice showing reduced ribbon synapses onto inner hair cells (blue) compared to sham-exposed mice. The CtBP2 (red) and GluR2 (green) are pre- and postsynaptic markers, respectively. **(i)** Average ribbon synapses onto per inner hair across the tonotopic region of the cochlea from noise-exposed and sham-exposed mice. (Noise vs. sham, 2-way ANOVA; exposure x frequency, F = 24.2, p = 1.1 × 10^−9^; effect of exposure, F = 126.3, p < 1.9 × 10^−10^; *, p < .05, compared to sham-exposed, Holm-Bonferroni’s post hoc). **(k)** Schematic of operant auditory avoidance task. **(l)** Plots of average sound-detection performance against the sound levels tested 10 days after noise (n = 5) and sham (n = 4) exposure. **(m)** Bar graph representing the average ABR and perceptual thresholds from noise- and sham-exposed mice. Filled circles represent individual data point. (Kruskal–Wallis test: Kruskal-Wallis statistic = 13.63, p < 1.1 × 10^−4^; *, p < .05, Holm-Bonferroni’s post hoc).

To test for perceptual recovery, we employed an operant auditory avoidance task (**Fig. 1k**). Namely, following a 6-sec long noise-bursts at 70 dB SPL, mice were trained to cross from one side of the shuttle-box to another side to avoid a mild foot-shock (200 - 400 μA) (Methods). A successful crossing during the noise-bursts trial was called *Hit*, whereas crossing during a random 6-sec long silent window was called *False-Alarm (FA)* (**Fig. 1k**). Once mice completed their behavioral training, we measured their perceptual sound thresholds (**Fig.1l**). To do this, we presented noise-bursts at various sound intensity levels (20-80 dB SPL) in random order, measured the Hit and FA rates at individual sound-levels, and then calculated sound-detection rate (Hit% - FA%). To quantify the perceptual sound detection threshold, we plotted the sound-detection rate against the sound-level and the sound-level with a 50% sound-detection rate was defined as the sound-detection threshold (**Fig. 1l**). When we measured the detection thresholds ten days after noise trauma, we found that perceptual thresholds in noise-exposed mice were significantly lower than ABR thresholds, and almost identical to the sham-exposed mice’s perceptual threshold (**Fig. 1m**). These results support that ten days after peripheral damage perceptual hearing thresholds have fully recovered, despite the persistent peripheral damage as evidenced by increased peripheral hearing thresholds, and thus suggest the involvement of central plasticity mechanisms in this recovery. Given the full recovery of perceptual thresholds within ten days after noise trauma, we opted to study the mechanisms of cortical plasticity for ten days after trauma.

To study the mechanisms of central plasticity after peripheral damage, we focused on the primary auditory cortex (A1), which is a site of robust plasticity after peripheral damage^5-7,10,11,13,14,26,55-59^. Because the cellular mechanisms of this plasticity are not fully understood, we investigated the plasticity in the different neuronal subtypes. We first investigated the plasticity in the sound-evoked activity of A1 principal neurons (PNs) one, three, and ten days after noise trauma (**Fig. 2**). To selectively image sound-evoked responses from populations of PNs, we used adeno-associated virus (AAV) driven by the calcium/calmodulin-dependent protein kinase 2 (CaMKII) promoter to express the genetically encoded calcium indicator GCaMP6f (AAV-CaMKII-GCaMP6f) in putative PNs (**Fig. 2ab**). Twelve to 16 days after stereotaxic viral injections of GCaMP6f (**Fig. 2a**), we employed acute *in vivo* wide-field transcranial fluorescent imaging in head-fixed unanesthetized (awake) mice (**Fig. 2b**). After localizing A1 (Methods), we presented broadband sounds (6-64 kHz, 100 ms long) at 30-80 dB SPL and imaged the sound-evoked changes in the A1 GCaMP6f fluorescence (ΔF/F%) (**Fig. 2c**). Each sound was presented 8-10 times in a pseudo-random order. We first measured PNs’ response threshold, the sound-level which elicits a significant response. Consistent with the increased ABR wave I response threshold, identified as the AN threshold, we found that the PNs’ response threshold was significantly increased 1 and 3 days after NIHL (**Fig. 2d**). However, 10 days after NIHL, PNs’ response threshold was significantly lower than the AN threshold (**Fig. 2d** right). Importantly, when we compared the PNs’ response threshold on day 10 after NIHL with the perceptual threshold on day 10 after NIHL (**Fig. 2d** right), we did not find a significant difference, suggesting that the reduced response threshold A1 PNs after NIHL may contribute to, or at least is consistent with, the recovery of the perceptual threshold after peripheral damage.

**Figure 2.**
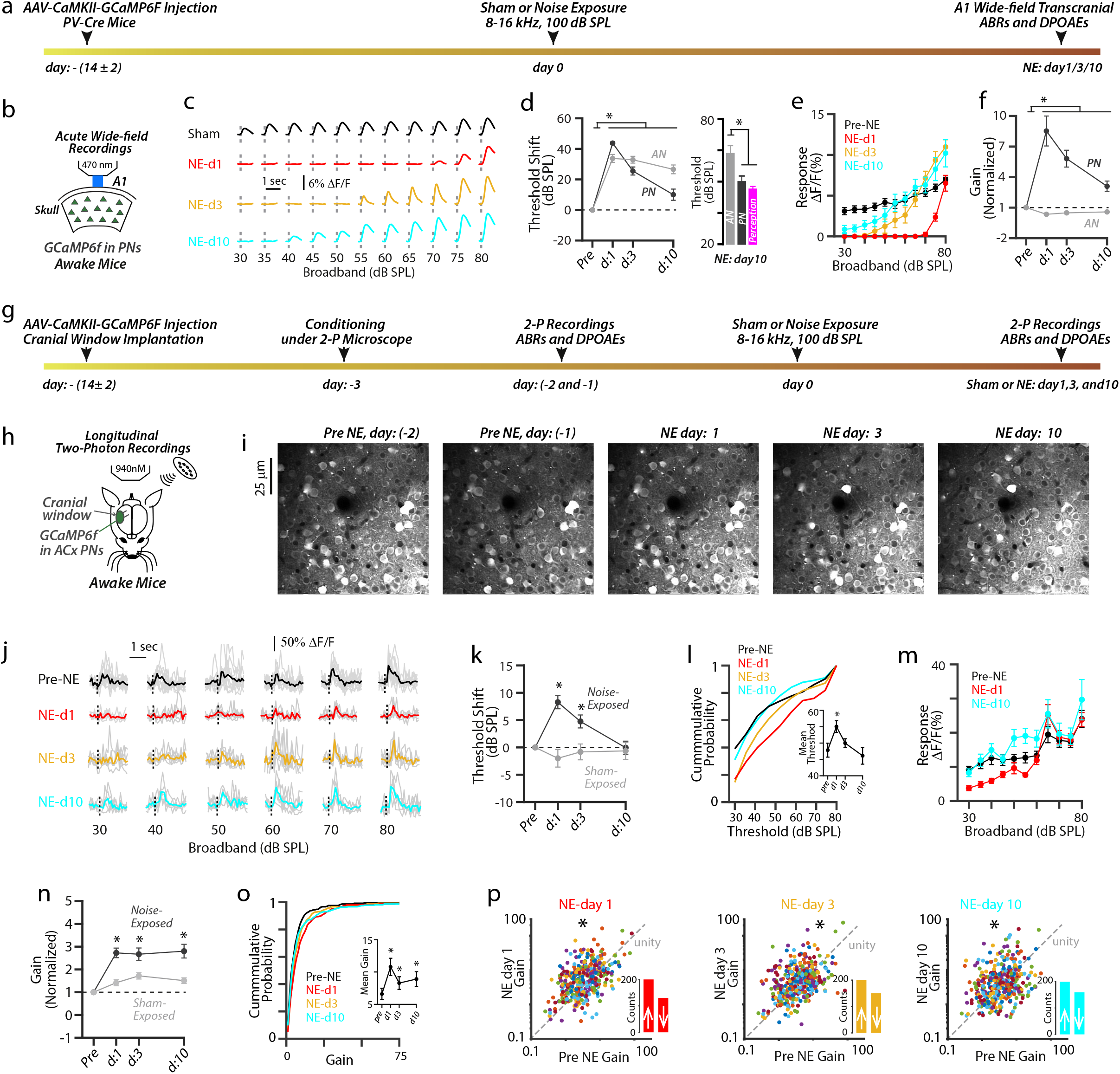
Robust sound-evoked activity (recovery) of A1 L2/3 PN neurons after NIHL. **(a)** Timetable of wide-field (WF) imaging experimental design for A1 PNs. **(b)** Schematic of experimental setup illustrating transcranial imaging of A1 PNs using GCaMP6f in a head-fixed awake mouse. **(c)** Representative transcranial fluorescence responses of A1 PNs to broadband sounds from sham- and noise-exposed mice. **(d)** Left: Average change in response thresholds of A1 PNs (dark grey) at 1, 3, and 10 days after noise exposure. (Noise: 8-11 mice vs. sham: 3 mice, mixed model ANOVA; exposure x time interaction, F = 30.4, p = 4.3 × 10^−9^; effect of exposure, F = 63.5, p = 3.8 × 10^−6^; *, p < 0.05, compared to pre-noise-exposed, Holm-Bonferroni’s post hoc). Average change in AN threshold (light grey) reproduced from **Figure 1**. Right: Bar graphs representing the average PNs, AN, and perception thresholds. (1-way ANOVA; F = 8.6, p = 6.8 × 10^−4^; *, p < 0.05, compared to AN, Holm-Bonferroni’s post hoc). **(e)** Average sound-evoked responses of A1 PNs to broadband sounds from noise-exposed mice. (n = 8-11 mice, 2-way ANOVA; time x sound level interaction, F = 5.06, p < 1.1 × 10^−10^; effect of time, F = 66.18, p = < 1.1 × 10^−10^; Compared to pre-noise-exposed: NE-day1, p = 2.8 × 10^−5^; NE-day3, p = 0.0007 and NE-day10, p > 0.99, Holm-Bonferroni’s post hoc; Pre-NE vs. NE-day10 responses to 75 and 80 dB SPL, p < 0.05, Holm-Bonferroni’s post hoc). **(f)** Average response gain of A1 PNs (dark grey) normalized to pre-noise-exposed gain after noise exposure at 1, 3, and 10 days. (Noise vs. sham, mixed model ANOVA; exposure x time interaction, F = 30.4, p = 4.3 × 10^−10^; effect of exposure, F = 63.5, p = 3.8 × 10^−6^; *, p < .05, compared to pre-noise-exposed, Holm-Bonferroni’s post hoc). Normalized AN gain (light grey) reproduced from **Figure 1. (g)** Timetable of longitudinal 2-photon imaging experimental design for A1 L2/3 PNs. **(h)** Schematic of experimental setup illustrating 2-photon imaging of A1 L2/3 PNs via cranial glass windows. **(i)** Z-stack images of a population of A1 L2/3 PNs tracked before and after NIHL. **(j)** Representative sound-evoked responses from an A1 L2/3 PN before and after NIHL. **(k)** Average change in response threshold of A1 L2/3 individual PNs from noise (dark grey) and sham (light grey) exposed mice. (Noise-exposed: 358 neurons from 11 mice, sham-exposed: 218 neurons from 5 mice, noise vs. sham: 2-way ANOVA; exposure x time interaction, F = 12.4, p = 5.0 × 10^−8^; effect of exposure, F = 11.6, p = 6.9 × 10^−4^; *, p < .05, compared to pre-noise-exposed, Holm-Bonferroni’s post hoc). **(l)** Cumulative response threshold of A1 L2/3 PNs, before and after NIHL. Inset: Average mean threshold of PNs per mouse (Friedman test; friedman statistic = 11.95, p = 0.007. *, p < .05, compared to pre-noise-exposed, Holm-Bonferroni’s post hoc). **(m)** Average sound-evoked responses of A1 L2/3 individual PNs to broadband sounds from noise-exposed mice. (2-way ANOVA; effect of time, F = 12.55, p = 5.3 × 10^−7^; compared to pre-noise-exposed, NEday1: p = 0.01, NEday3: p = 0.80, and NEday10: p = 0.001; Holm-Bonferroni’s post hoc). **(n)** Average gain of A1 L2/3 individual PNs normalized to pre-exposed gain from noise (dark grey) and sham (light grey) mice. (Noise vs. sham: 2-way ANOVA; exposure x time interaction, F = 4.7, p = 0.002; effect of exposure, F = 23.3, p = 1.7 × 10^−6^; *, p < .05, compared to pre-noise-exposed, Holm-Bonferroni’s post hoc). **(o)** Cumulative gain of A1 L2/3 PNs, before and after NIHL. Inset: Average mean gain of PNs per mouse (Friedman test; friedman statistic = 10.31, p = 0.01. *, p < .05, compared to pre-noise-exposed, Holm-Bonferroni’s post hoc). **(p)** Scatter plots of the gain of individual A1 L2/3 PNs before and after NIHL. Dotted line represents unity. Insets: Bar graphs representing the number of neurons showing increased gain (↑ above unity) and reduced gain (↓ below unity) after NIHL. PreNE vs. NEday1: p = 9.9 ×10^−5^, PreNE vs. NEday3: p = 0.006, and PreNE vs. NEday10: p = 0.007; permutation test.

Next, we measured the amplitudes of sound-evoked responses of A1 PNs (**Fig. 2e**). We found that PN response amplitudes were reduced 1 day after NIHL (**Fig. 2e**, red), but showed significant recovery in 3 and 10 days after NIHL (**Fig. 2e**, cyan), and even surpassed pre-noise-exposed response amplitudes in response to suprathreshold sound levels (at 75 and 80 dB SPL). We next quantified the response gain of sound-evoked activity of A1 PNs (**Fig. 2f**). In contrast to ABR wave I response gain (AN gain) which remained decreased after noise trauma (**Fig. 2f**, light grey), PN gain was increased and remained increased elevated during the 10 days after NIHL (**Fig. 2f**, dark grey), which is consistent with previous results^10,16,17^. Moreover, we did not find any changes in response threshold, response amplitude, and gain in sham-exposed mice (**supplement Fig. 2b-d**). Together, these results suggest that ten days after peripheral damage, A1 PNs display increased gain and recovered response thresholds and amplitudes.

Wide-field imaging reflects neuronal responses arising from different neuronal compartments (e.g., somata, dendrites, and axons) and different cortical layers^60^. Moreover, wide-field imaging reflects responses from a population of neurons, but individual neurons may have distinct sound-evoked responses (e.g., recovered vs. non-recovered) after NIHL. To address these caveats and questions, we performed longitudinal two-photon imaging of the same individual A1 L2/3 PNs in awake mice for 10 days after NIHL (**Fig. 2g-p, and supplement Fig. 2e-j**). After locating A1, we presented trains of broadband sounds and imaged the sound-evoked responses of individual A1 L2/3 PNs’ somata (**Fig. 2i, j**). To use each neuron as its own control, we tracked the same individual neurons for 10 days after NIHL (**Fig. 2i**). Pre-exposure sessions lasted two days, and average responses of individual neurons from both days were used as pre-exposure responses. After motion and neuropil correction (Methods), we were able to track 531 L2/3 PNs from 11 mice for 10 days after NIHL. To identify the sound-responsive neurons, we first calculated the individual neurons’ tuning strength during pre-exposure conditions, and only the neurons with d’ ≥ 0 were analyzed further (n = 358/531 PNs from 11 mice, Methods). Consistent with our transcranial results, we found that the response thresholds of individual L2/3 PNs’ were fully recovered 10 days after NIHL and had a similar cumulative distribution of response thresholds compared to pre-noise-exposure thresholds (**Fig. 2k, l**). Also, we found that the sound-evoked responses of individual PNs were reduced 1 day after NIHL, but overall recovered or surpassed pre-noise-exposure responses 10 days after NIHL (**Fig. 2m**). Also, consistent with our transcranial results, the gain of individual PNs’ was increased after NIHL and remained elevated even 10 days after NIHL (**Fig. 2n**), showing a shift in cumulative distribution towards higher gain (**Fig. 2o**). When we plotted individual PN gain after noise-exposure against pre-noise-exposure gain, we also found that on average the gain was increased after noise exposure (**Fig. 2p**) and the majority of PNs showed increased gain after NIHL (**Fig. 2p insets and supplement Fig. 2j**: day1: 228/358, day3: 208/358, and day 10: 199/358). Finally, we did not observe a change in either the threshold (**Fig. 2k**) or the gain (**Fig. 2n**) in sham-exposed mice (218 neurons from 5 mice, **Fig. 2k, n and supplement Fig. 2f-j)**. Together, our results show that despite peripheral damage, A1 L2/3 PNs show recovered response thresholds, response amplitudes, and increased response gain. The central goal of this study is to identify the core mechanism underlying this PN recovery from peripheral damage.

To begin, we considered the possible role of inhibitory circuitry on the recovery of A1 L2/3 PNs after NIHL. To this end, we first used a computational model to investigate the possible changes in inhibition that can achieve PN high gain. Past modeling work has shown that a decrease in the recurrent inhibition in a recurrently coupled cortical model results in higher PN gain^53,61^, thus an NIHL-induced reduction in inhibition could be a candidate mechanism. However, strong recurrent PN connections can yield unstable, runaway behavior if a recurrently coupled inhibitory population is unable to dynamically track and cancel the recurrent excitatory activity^51,53,62^. As such, the stabilization role for inhibition poses constraints on how reduced inhibition can safely modulate cortical gain, because a global and non-specific inhibitory reduction could lead to instability and pathology, such as epileptic-like activity^52^. Thus, a simplified two-population model, consisting of generic excitatory and inhibitory neurons, would likely fall short of capturing the experimental results presented thus far^53^ (**Fig. 2**). As a result, we started our investigation by considering a computational network of leaky integrate-and-fire neuron models (Methods) of three subpopulations of neurons (PN, PV, and SOM neurons) (**Fig. 3a**). PN and PV neurons received a feedforward presynaptic drive, and we modeled sound level by increasing the firing rate of the feedforward inputs. We considered four sound levels: none (no sound), low, medium, and high. The control (pre-exposure) behavior of the network lies in an asynchronous (stable) regime, with the firing rate of all three populations increasing monotonically with sound level (**Fig. 3b, c**). Since peripheral damage reduces the intensity of peripheral sensory drive from the cochlea to the AN and the brain, noise-induced damage in our model is implemented by decreasing the feedforward (evoked) and background (spontaneous) firing rates. We modeled recovery after NIHL, as observed 10 days after NIHL (**Fig. 2**), either as a static depolarization or hyperpolarization of individual cortical neurons. The underlying cause behind these inputs could be due to intrinsic or synaptic mechanisms that restore neuronal threshold post NIHL. Consistent with our prediction on the constraints on how reduced inhibition can safely modulate cortical gain, we found that depending on the magnitude and sign of these currents to each subpopulation, the network spiking behavior varied drastically, from oscillatory and unstable, to asynchronous and stable with high gain (**Fig. 3d**).

**Figure 3.**
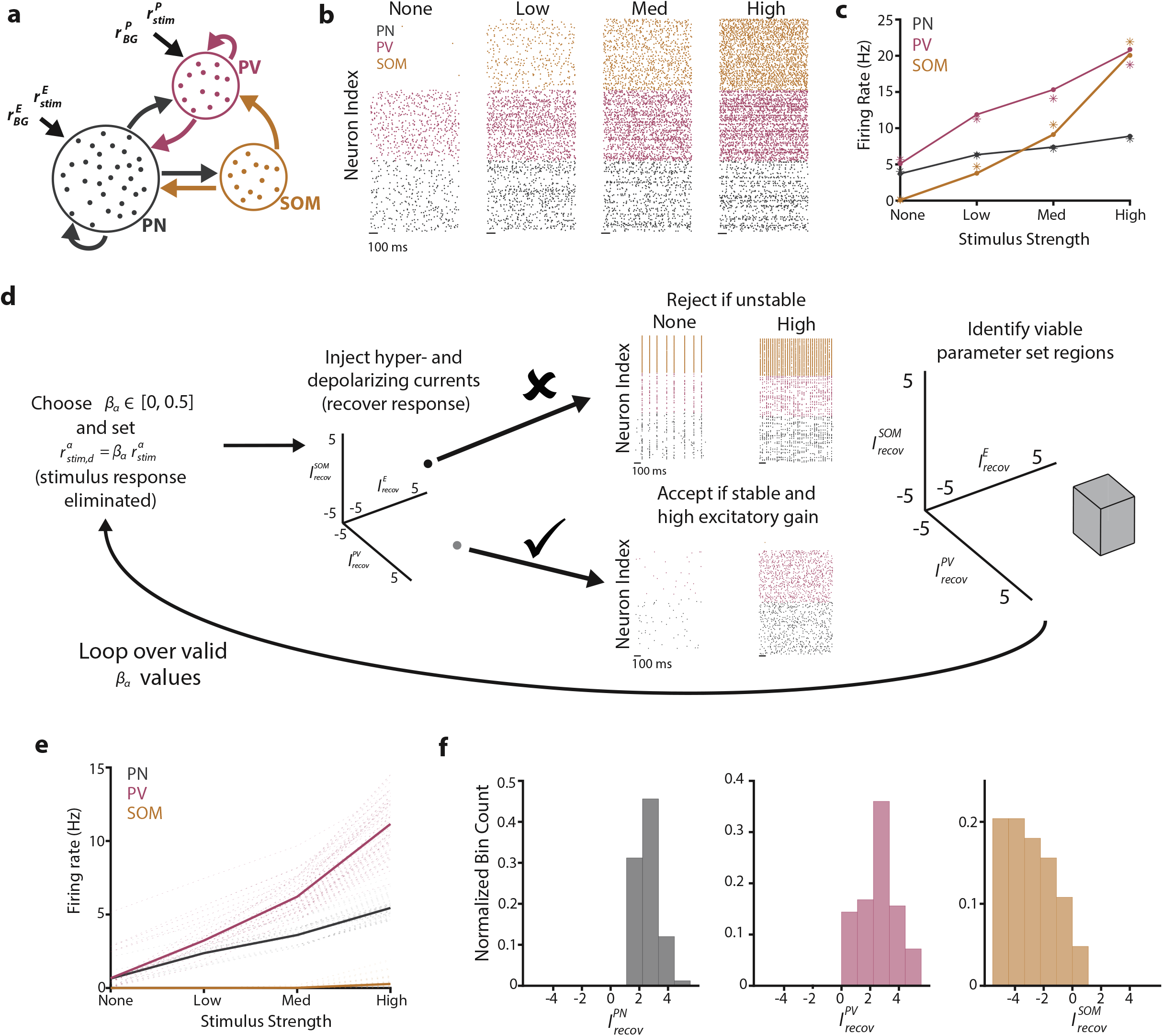
Three-population model suggests that SOM suppression is responsible for threshold and gain recovery. **(a)** Schematic of connectivity across the three populations (PN, SOM, and PV). **(b)** Raster plots showing the spiking activity of a subset of neurons at four stimulus levels for the PN (black), PV (magenta), and SOM populations (orange). **(c)** Firing rate for the spiking model (dot-line) and mean-field theory (asterisks). **(d)** Schematic of the parameter sweep algorithm. For specific pairs of damage values (*β*_*PN*_, *β*_*PN*_), the mean-field theory was used to find the firing rates of the model for points in the recovery current space 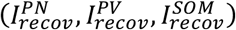. Parameter sets that yielded stable behavior (asynchronous), along with a low threshold and improved gain (bottom arrow) were accepted, while all others were rejected (e.g., oscillatory, top arrow). Viable parameter regions were identified. This process was looped over all damage values. **(e)** Firing rate for the three populations. Translucent lines correspond to distinct parameter sets, while bolded lines are the average firing rates across all viable parameter sets. **(f)** Histograms of the recovery currents were found in the viable parameter sets for the three populations. All parameter values can be found in Tables 1 and 2.

Because our major focus is to understand the circuit pathways responsible for the recovery of PNs’ threshold, high gain, and stability after NIHL, we utilized a mean-field circuit theory (see Methods), which captures the average neuronal firing rate for each of the subpopulations, to perform an extensive, brute force parameter sweep (**Fig. 3c, d**). Viable parameter sets that matched our experimental observations of PNs were defined as those that produced stable network dynamics with lower PN response thresholds and higher PN gain than in control (see Methods for additional details). Parameter sets that met these criteria yielded average SOM firing rates that were suppressed compared to control, exhibiting little-to-no SOM recovery after damage and PV neurons with recovery similar to that of the PN population (**Fig. 3e**). These successful parameter sets can be further explored by examining the strength and sign (depolarizing vs. hyperpolarizing) of the recovery currents injected into each of the subpopulations (**Fig. 3f**). Specifically, while the PN and PV neurons received depolarizing inputs, SOM cells largely received hyperpolarizing inputs. These results suggest that the selective suppression of SOM neurons allows for PNs to overcome the loss of feedforward input and thereby recover their response threshold and exhibit higher gain after peripheral damage compared to the control (sham-exposed) case. These modeling results lead to a pair of testable hypotheses: 1) the PV population will have a matched recovery to that of the PN populations, and 2) SOM neurons will not recover post-NIHL.

**Table 1:** Default parameter values.

| Parameter | Value | Description |
| --- | --- | --- |
| $\tau_m$ | 10 (ms) | membrane time constant |
| $\tau_s$ | 0.5 (ms) | synaptic time constant |
| $\tau_r$ | 2 (ms) | refractory period |
| $E_L$ | -65 (mV) | resting potential |
| $V_{th}$ | -50 (mV) | spike threshold |
| $V_r$ | -65 (mV) | reset potential |
| $w$ | 0.6 (mV) | synaptic strength of the excitatory connection |
| $g$ | 3 | the synaptic factor for inhibitory connection |
| $N_e$ | 5000 | Num. of PN neurons |
| $N_p$ | 520 | Num. of PV neurons |
| $N_s$ | 520 | Num. SOM neurons |
| $N_v$ | 0 / 520 | Num. VIP neurons (3/4 populations) |
| $N_{ext}^e$ | 500 | Num. external inputs to PN neurons |
| $N_{ext}^p$ | 400 | Num. external inputs to PV neurons |
| $N_{ext}^s$ | 0 | Num. external inputs to SOM neurons |
| $N_{ext}^v$ | -- / 400 | Num. external inputs to VIP neurons (3/4 populations) |
| $\sigma_{fixed}^2$ | 10 | Fixed background noise level |
| $r_{bg}^a$ | 3 (Hz) | Background excitatory firing rate |
| $r_{stim}^a$ | 0, 2, 4, or 8 (Hz) | Stimulus firing rate (none, low, med, high) |
| $\gamma$ | 0.5 | Damage to background firing rate |
| $\beta^a$ | [0.05, 0.5] | Damage to input stimulus firing rate to pop $a$ |
| $I_{recov}^a$ | [-5, 5] | Recovery current to pop $a$ |
| $\kappa_{thres}$ | -0.7 | Stability threshold |

**Table 2:** The probability of a connection between presynaptic (columns) and postsynaptic (rows) populations.

|  | E | PV | SOM | VIP |
| --- | --- | --- | --- | --- |
| E | 0.03 | 0.10 | 0.10 | 0 |
| PV | 0.05 | 0.10 | 0.07 | 0 |
| SOM | 0.05 | 0 | 0 | 0.10 |
| VIP | 0.05 | 0.15 | 0.05 | 0 |

To test the first hypothesis, we first investigated the effect of NIHL on response threshold, amplitude, and gain in A1 L2/3 PV neurons. To selectively target and image sound-evoked responses from populations of PV neurons, we injected AAV expressing Cre-dependent GCaMP6f (AAV-Flex-GCaMP6f) into the A1 of PV-Cre mice **(Fig. 4a)**. We first employed *in vivo* wide-field transcranial imaging of populations of PV neurons in awake mice (**Fig. 4a-f**). We found that the response threshold of PV neurons was increased 1 day after noise exposure (**Fig. 4d**, magenta). However, 3 days after noise exposure, the PV population response threshold was lower than the response threshold of A1 PNs (**Fig. 4d**), suggesting that the response thresholds of PV neurons are recovered even before the response threshold of PNs. Ten days after NIHL, PV neurons response thresholds remain low and were not different from the PN response thresholds (**Fig. 4d**). Moreover, we found the reduced sound-evoked response amplitudes of A1 PV neurons 1 day after NIHL (**Fig. 4e**, red), which recovered by 10 days after NIHL **(Fig. 4e**, cyan). Importantly, we found that noise exposure increased the gain of the PV population, which remained elevated for 10 days after NIHL (**Fig. 4f**). We did not observe a change in the PV population response threshold and gain in sham-exposed mice (**supplement Fig. 4a, b**). Together, these results demonstrate that PV population recovery is overall similar to PN recovery.

**Figure 4.**
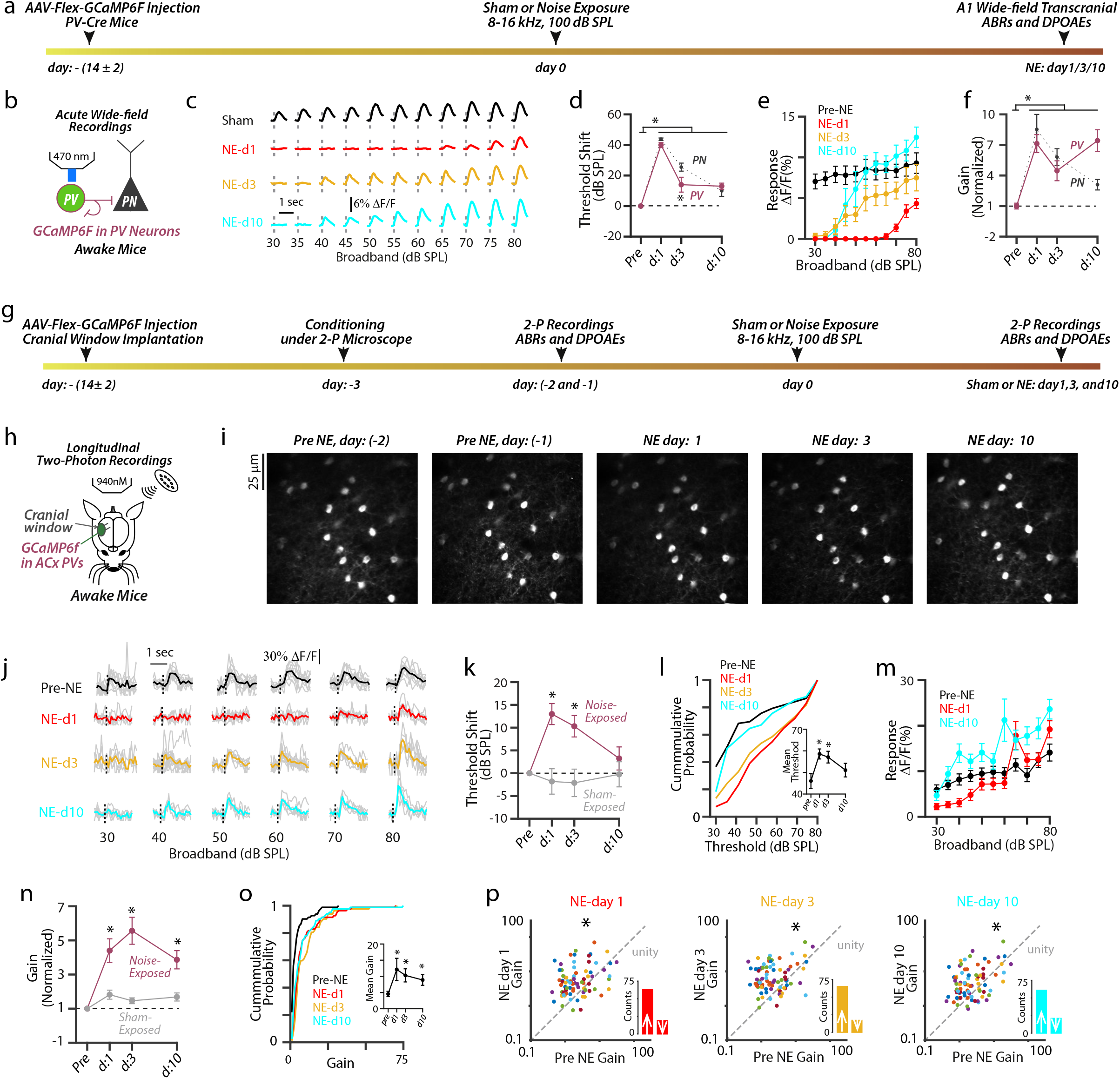
Robust sound-evoked activity (recovery) of A1 L2/3 PV neurons after NIHL. **(a)** Timetable of wide-field (WF) imaging experimental design for A1 PV neurons. **(b)** Schematic of the cortical circuit and experimental setup illustrating transcranial imaging of A1 PV neurons using GCaMP6f in a head-fixed awake mouse. **(c)** Representative transcranial fluorescence responses of A1 PV neurons to broadband sounds from sham- and noise-exposed mice. **(d)** Average change in response thresholds of A1 PV neurons (magenta) at 1, 3, and 10 days after noise exposure. (Noise: 5-6 mice vs. sham: 3 mice, mixed model ANOVA; exposure x time interaction, F = 22.4, p = 2.8 × 10^−7^; effect of exposure, F = 89.0, p = 1.0 × 10^−9^; *, p < 0.05, compared to pre-noise-exposed, Holm-Bonferroni’s post hoc). Average change in PN threshold (dark grey) reproduced from **Figure 2**. (PV vs. PN: mixed model ANOVA; cell-type x time interaction, F = 3.7, p = 0.01; effect of time, F = 120, p < 10^−10^; *, p < 0.05, compared to PNs at NEday3, Holm-Bonferroni’s post hoc). **(e)** Average sound-evoked threshold responses of A1 PV neurons to broadband sounds from noise-exposed mice. (n = 5-6 mice, 2-way ANOVA; time x sound level interaction, F = 2.9, p = 3.3 × 10^−6^; effect of time, F = 98.5, p < 1.1 × 10^−10^; Compared to pre-noise-exposed: NE-day1, p < 1.1 × 10^−10^; NE-day3, p = 4.5 × 10^−10^, and NE-day10, p = 0.11, Holm-Bonferroni’s post hoc). **(f)** Average response gain of A1 PV neurons (magenta) normalized to pre-noise exposed gain after noise exposure at 1, 3, and 10 days. (Noise vs. sham, mixed model ANOVA; exposure x time interaction, F = 7.4, p = 9.6 × 10^−4^; effect of exposure, F = 53.7, p = 1.1 × 10^−7^; *, p < .05, compared to pre-noise-exposed, Holm-Bonferroni’s post hoc). Normalized PN gain (dark grey) reproduced from **Figure 2. (g)** Timetable of longitudinal 2-photon imaging experimental design for A1 L2/3 PV neurons. **(h)** Schematic of experimental setup illustrating longitudinal 2-photon imaging of A1 L2/3 PV neurons. **(i)** Z-stack images of a population of A1 L2/3 PV neurons tracked before and after NIHL. **(j)** Representative sound-evoked responses from an A1 L2/3 PV before and after NIHL. **(k)** Average change in response threshold of A1 L2/3 individual PV neurons from noise (magenta) and sham (grey) exposed mice. (Noise-exposed: 82 neurons from 6 mice, sham-exposed: 80 neurons from 7 mice, noise vs. sham: 2-way ANOVA; exposure x time interaction, F = 7.3, p = 8.5 × 10^−5^; effect of exposure, F = 11.17, p = 0.001; *, p < .05, compared to pre-noise-exposed, Holm-Bonferroni’s post hoc). **(l)** Cumulative response threshold of A1 L2/3 PV neurons, before and after NIHL. Inset: Average mean threshold of PV neurons per mouse (Friedman test; friedman statistic = 11.53, p = 0.003. *, p < .05, compared to pre-noise-exposed, Holm-Bonferroni’s post hoc). **(m)** Average sound-evoked responses of A1 L2/3 individual PV neurons to broadband sounds from noise-exposed mice. (2-way ANOVA; time x sound level interaction, F = 2.2, p = 3.4 × 10^−4^; effect of time, F = 29.9, p < 1.1 × 10^−10^; Compared to pre-noise-exposed: NE-day1, p > 0.99; NE-day3, p = 1.7 × 10^−7^ and NE-day10, p < 1.1 × 10^−10^, Holm-Bonferroni’s post hoc. Compared to pre-noise-exposed 50 dB SPL and lower: NE-day1, p <0.05, Holm-Bonferroni’s post hoc). **(n)** Average gain of A1 L2/3 individual PV neurons normalized to pre-exposed gain from noise (magenta) and sham (grey) mice. (Noise vs. sham: 2-way ANOVA; exposure x time interaction, F = 10.1, p = 1.6 × 10^−6^; effect of exposure, F = 26.9, p = 6.1 × 10^−7^; *, p < .05, compared to pre-noise-exposed, Holm-Bonferroni’s post hoc). **(o)** Cumulative gain of A1 L2/3 PV neurons, before and after NIHL. Inset: Average mean gain of PV neurons per mouse (Friedman test; friedman statistic = 12.2, p = 0.002. *, p < .05, compared to pre-noise-exposed, Holm-Bonferroni’s post hoc). **(p)** Scatter plots of the gain of individual A1 L2/3 PV neurons before and after NIHL. Dotted line represents unity. Insets: Bar graphs representing the number of neurons showing increased gain (↑ above unity) and reduced gain (↓ below unity) after NIHL. PreNE vs. NEday1: p = 9.9 ×10^−5^, PreNE vs. NEday3: p = 0.006, and PreNE vs. NEday10: p = 0.007; permutation test. PreNE vs. NEday1: p = 3.9 ×10^−4^, PreNE vs. NEday3: p = 9.9 ×10^−5^, and PreNE vs. NEday10: p = 5.9 ×10^−4^; permutation test.

Next, we performed longitudinal 2P imaging of A1 L2/3 PV neurons (**Fig 4g-j**). We tracked and included in our analysis 82 PV neurons from 6 mice for 10 days after NIHL. Consistent with our transcranial imaging results, PV neurons displayed recovered response thresholds (**Fig. 4k, l**), and even surpassed pre-noise-exposure responses 10 days after NIHL amplitudes (**Fig. 4m, cyan**). Moreover, the gain of individual PV neurons increased after NIHL and remained increased during the 10 days after NIHL (**Fig. 4n-p**). The majority of PV neurons showed increased response gain after NIHL (day1: 63/82, day3: 65/82, and day10: 61/82) (**Fig. 4p insets and supplement Fig. 4g**). We did not observe any changes in the response threshold, amplitude, and gain of L2/3 PV neurons in sham-exposed mice (80 neurons from 7 mice, **Fig. 4k, n and supplement Fig. 4c-g**). These results demonstrate that, in response to peripheral damage, A1 L2/3 PV neurons match the recovery of PNs to act as stabilizers of A1 network activity, validating the first modeling hypothesis. Consequently, PV neurons likely do not contribute to the increased PN gain after recovery from NIHL.

To test our second hypothesis, we investigated the role of SOM neurons during recovery from NIHL. We started our investigation with *in vivo* wide-field transcranial imaging (**Fig. 5**) and found that the response threshold of the A1 SOM population was very high 1 day after NIHL, above 80 dB (**Fig. 5c**, red). Importantly, response thresholds did not recover and remained significantly higher than the response threshold of PV neurons and PNs even 10 days after noise exposure (**Fig. 5d**). Additionally, response amplitudes were reduced after NIHL and did not fully recover even 10 days after NIHL (**Fig. 5e**, cyan). Finally, we did not observe any gain changes in SOM population response (**Fig. 5f**). We did not observe a change in the response threshold and gain of SOM neurons in sham-exposed mice (**supplement Fig. 5a-b)**. Overall, in contrast to the robust sound-evoked PN and PV neurons activity after noise trauma, SOM neurons’ sound-evoked activity remained significantly reduced throughout the 10 days after noise trauma.

**Figure 5.**
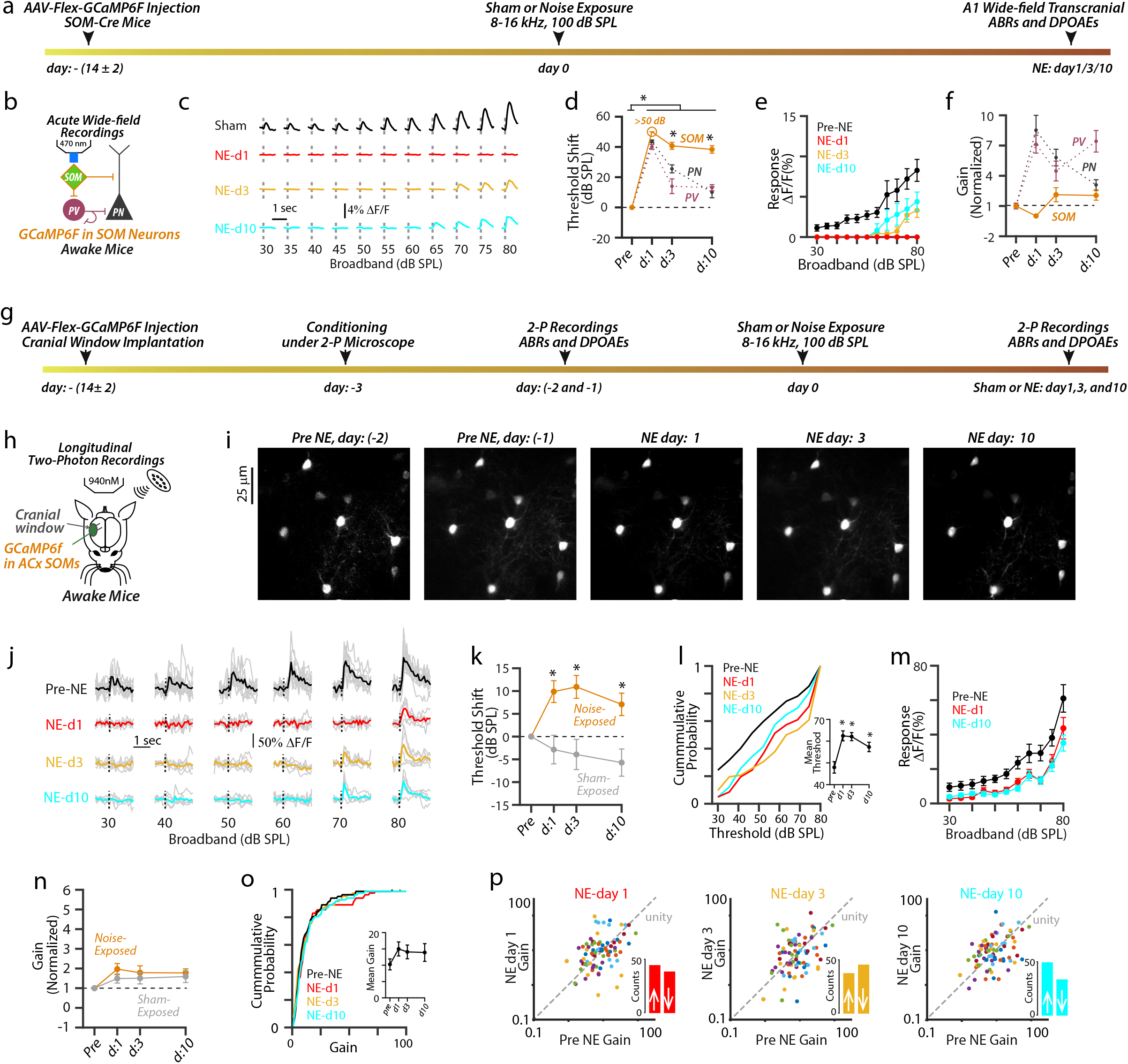
Reduced sound-evoked activity (recovery) of A1 L2/3 SOM neurons after NIHL. **(a)** Timetable of wide-field (WF) imaging experimental design for A1 SOM neurons. **(b)** Schematic of the cortical circuit and experimental setup illustrating transcranial imaging of A1 SOM neurons using GCaMP6f in a head-fixed awake mouse. **(c)** Representative transcranial fluorescence responses of A1 SOM neurons to broadband sounds from sham- and noise-exposed mice. **(d)** Average change in response thresholds of A1 SOM neurons (orange) after noise exposure at 1, 3, and 10 days. Note: Since we did not observe an sound-evoked activity in SOM neurons at 1 day after noise-exposure even at 80 dB SPL sounds, we did not assign a threshold shift at this time point, but it is >50 dB SPL (Noise: 5-6 mice vs. sham: 4 mice, mixed model ANOVA; exposure x time interaction, F = 81.4, p < 10^−10^; effect of exposure, F = 682.1, p < 10^−10^; *, p < 0.05, compared to pre-noise-exposed, Holm-Bonferroni’s post hoc). Average change in PN (grey) and PV (red) threshold reproduced from **Figure 4**. (PV vs. PN vs. SOM neurons: mixed model ANOVA; cell-type x time interaction, F = 11.43, p = 7.65 × 10^−8^; effect of time, F = 181.6, p < 10^−10^; *, p < 0.05, compared to PNs and PV neurons at NEday3 and NEday10, Holm-Bonferroni’s post hoc). **(e)** Average sound-evoked responses of A1 SOM neurons to broadband sounds from noise-exposed mice. (n = 5-6, 2-way ANOVA; time x sound level interaction, F = 2.8, p = 2.3 × 10^−5^; effect of time, F = 82.97, p < 1.1 × 10^−10^; Compared to pre-noise-exposed: NE-day1, p < 1.1 × 10^−10^; NE-day3, p < 1.1 × 10^−10^ and NE-day10, p < 1.1 × 10^−10^, Holm-Bonferroni’s post hoc). **(f)** Average response gain of A1 SOM neurons (orange) normalized to pre-noise-exposed gain at 1, 3, and 10 days after noise exposure. (Noise: 5-6 mice vs. sham: 3 mice, mixed model ANOVA; exposure x time interaction, F = 2.1, p = 0.12; effect of exposure, F = 0.38, p = 0.54). Normalized PN (grey) and PV (red) neurons’ gain reproduced from **Figure 4. (g)** Timetable of longitudinal 2-photon imaging experimental design for A1 L2/3 SOM neurons. **(h)** Schematic of experimental setup illustrating longitudinal 2-photon imaging of A1 L2/3 SOM neurons. **(i)** Z-stack images of a population of A1 L2/3 SOM neurons tracked before and after NIHL. **(j)** Representative sound-evoked responses from A1 L2/3 SOM neurons before and after NIHL. **(k)** Average change in response threshold of A1 L2/3 individual SOM neurons from noise (orange) and sham (grey) exposed mice. (Noise-exposed: 82 neurons from 15 mice, sham-exposed: 42 neurons from 9 mice, noise vs. sham: 2-way ANOVA; exposure x time interaction, F = 5.3, p = 0.001; effect of exposure, F = 16.60, p = 8.2 × 10^−5^; *, p < .05, compared to pre-noise-exposed, Holm-Bonferroni’s post hoc). **(l)** Cumulative response threshold of A1 L2/3 SOM neurons, before and after NIHL. Inset: Average mean threshold of SOM neurons per mouse (Repeated measure one-way ANOVA; F = 11.02, p = 5.4 × 10^−5^. *, p < .05, compared to pre-noise-exposed, Holm-Bonferroni’s post hoc). **(m)** Average sound-evoked responses of A1 L2/3 individual SOM neurons to broadband sounds from noise-exposed mice. (2-way ANOVA; time x sound level interaction, F = 1.6, p = 0.02; effect of time, F = 45.97, p < 1.1 × 10^−10^; Compared to pre-noise-exposed: NE-day1, p = p < 1.1 × 10^−10^; NE-day3, p = p < 1.1 × 10^−10^ and NE-day10, p < 1.1 × 10^−10^, Holm-Bonferroni’s post hoc). **(n)** Average gain of A1 L2/3 individual SOM neurons normalized to pre-exposed gain from noise (orange) and sham (grey) mice. (Noise vs. sham: 2-way ANOVA; exposure x time interaction, F = 0.28, p = 0.83; effect of exposure, F = 1.2, p = 0.27). **(o)** Cumulative gain of A1 L2/3 SOM neurons, before and after NIHL. Inset: Average mean gain of SOM neurons per mouse (Repeated measure one-way ANOVA; F = 1.46, p = 0.24). **(p)** Scatter plots of the gain of individual A1 L2/3 SOM neurons before and after NIHL. Dotted line represents unity. Insets: Bar graphs representing the number of neurons showing increased gain (↑ above unity) and reduced gain (↓ below unity) after NIHL. PreNE vs. NEday1: p = 0.30, PreNE vs. NEday3: p = 0.71, and PreNE vs. NEday10: p = 0.36; permutation test.

Consistent with our transcranial results, longitudinal 2P imaging of individual A1 L2/3 SOM neurons (82 neurons from 15 mice) showed increased response thresholds after injury, which remained elevated throughout the 10 days after NIHL (**Fig. 5g-k**), and a shift in the cumulative distribution of response threshold towards higher sound levels (**Fig. 5l**). Also, consistent with the transcranial results, we found reduced sound-evoked amplitudes of individual SOM neuron response **(Fig. 5m, cyan)**. Moreover, we did not observe any change in the gain of A1 L2/3 SOM neurons after NIHL **(Fig. 5n-p)**. Finally, we did not observe a change in the SOM neurons’ response threshold, amplitude, and gain in sham-exposed mice (**Fig. 5k, n and supplement Fig. 5c-g**, 42 neurons from 9 mice). In total, these results support the second modeling hypothesis that SOM neurons’ responses are suppressed during recovery from NIHL. These results support the notion that the reduced activity of SOM neurons disinhibits PV neurons and PNs post-NIHL, thus allowing for high PV and PN response gain.

We next explored the mechanism underlying the SOM neuron suppression, which in turn contributes to the enhanced gain of L2/3 A1 PNs and PV neurons. The reduction in overall SOM activity might be due to changes in the intrinsic cellular makeup of SOM neurons, the synaptic input afferent to SOM neurons, or a combination of the two mechanisms. To test for changes in intrinsic properties, we performed *ex vivo* brain slice electrophysiology of AC L2/3 SOM neurons after NIHL **(Fig. 6)**. Due to the lack of cytoarchitectural features, it is challenging to locate the AC in brain slices. Therefore, to localize the AC, we labeled AC corticocollicular (CCol) L5B PNs (red) projecting to the inferior colliculus, by injecting red fluorescent retrograde microspheres into the inferior colliculus of SOM-GFP mice (**Fig. 6a**, Methods). The localization of CCol PNs in the AC (**Fig. 6b**), along with anatomical landmarks, such as the rhinal fissure and the underlying hippocampal formation allowed us to locate the AC as described previously^63-65^. After localizing the AC, we measured the intrinsic properties of AC L2/3 SOM neurons in noise- and sham-exposed mice (**Fig. 6b**). The input resistance (R_input_) and the membrane resting potential (V_rest_) did not change over the 10 days after noise-compared to sham-exposure **(Fig. 6c-e)**. Similarly, noise trauma did not affect action potential width (AP_width_), AP threshold (AP_threshold_), and firing rate of SOM neurons **(Fig. 6f-k)**. Finally, the firing rate adaptation ratio of the SOM neurons, calculated as the ratio of instantaneous firing frequency between the ninth and tenth AP and instantaneous frequency between the second and third AP (f9/f2)^64^, showed no significant difference between sham- vs. noise-exposed mice **(Fig. 6l, m)**. Taken together, these results suggest that the reduced sound-evoked activity in SOM neurons after NIHL is likely not due to changes in intrinsic, cellular properties in the SOM neurons themselves.

**Figure 6.**
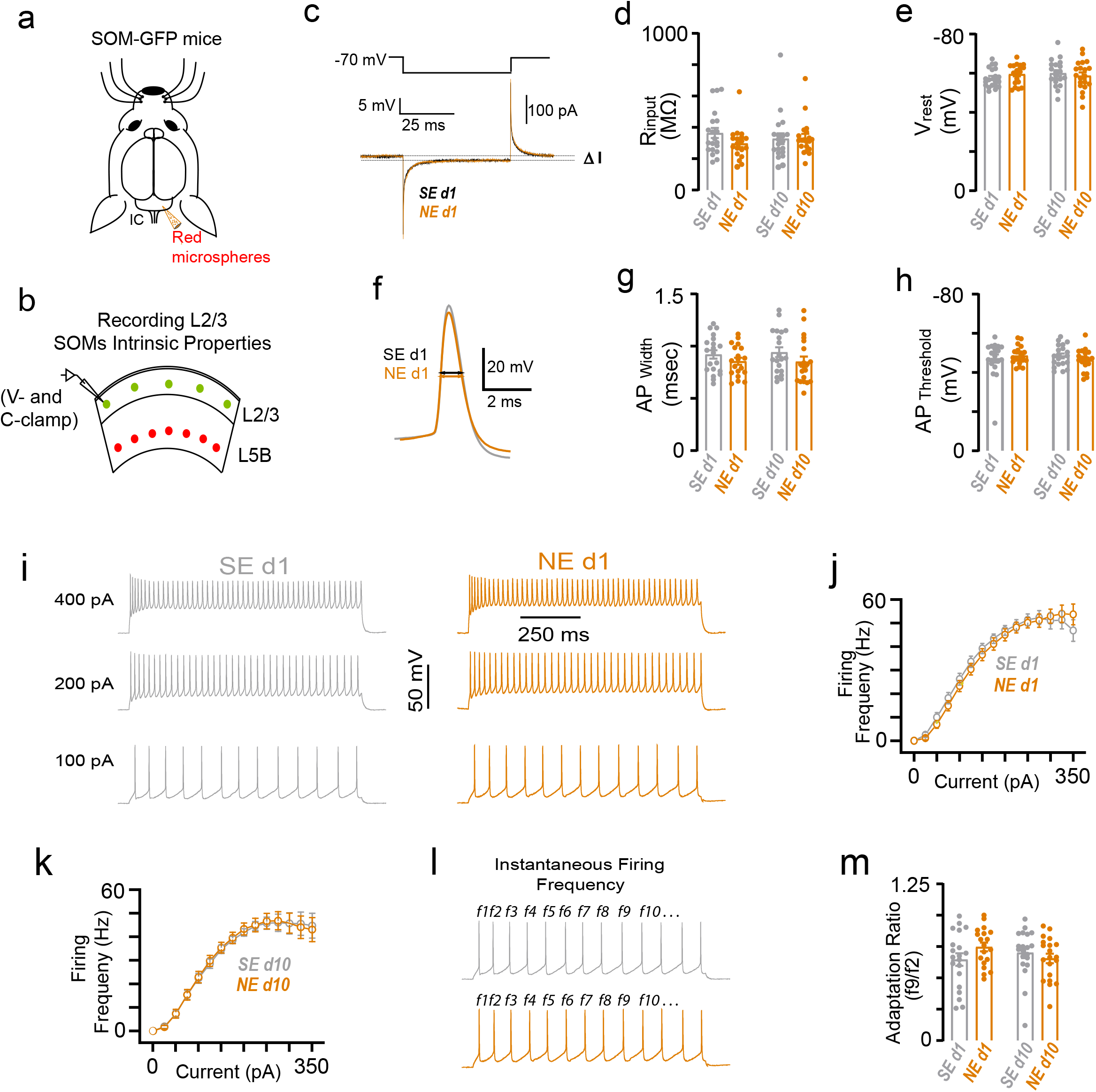
Intrinsic properties of AC L2/3 SOM neurons do not change after NIHL. **a)** Schematic illustration of stereotaxic injections of red retrograde microspheres into the right inferior colliculus to label corticocollicular neurons and identify the AC in the brain slices. **b)** Schematic illustration of brain slice electrophysiology experiment showing recordings of L2/3 SOM neurons’ intrinsic properties. Red circles represent the L5B corticocollicular neurons. Green circles represent the L2/3 SOM neurons. **c)** Schematic of hyperpolarizing pulses (top) and representative transient current (bottom) responses in SOM neurons in voltage-clamp recording mode. **d)** Average input resistance of L2/3 SOM neurons after noise- or sham-exposure. Filled circles represent the input resistance of individual SOM neurons. (SEday1: 20 neurons from 3 mice, NEday1: 19 neurons from 3 mice, SEday10: 20 neurons from 3 mice, and NEday10: 20 neurons from 3 mice, mixed model ANOVA; exposure x time interaction, F = 2.0, p = 0.16; effect of exposure, F = 0.89, p = 0.34). **e)** Average resting membrane potential of SOM neurons after noise- or sham-exposure. Filled circles represent the resting membrane potential of individual SOM neurons. (Mixed model ANOVA; exposure x time interaction, F = 1.68, p = 0.20; effect of exposure, F = 0.07, p = 0.78). **f)** Representative action potential (AP) waveforms. Arrows indicate AP width. **g)** Average AP width of SOM neurons after noise- or sham-exposure. Filled circles represent AP width of individual SOM neurons. (Mixed model ANOVA; exposure x time interaction, F = 0.09, p = 0.76; effect of exposure, F = 2.6, p = 0.11). **h)** Average AP threshold of SOM neurons after noise- or sham-exposure. Filled circles represent AP threshold of individual SOM neurons. (Mixed model ANOVA; exposure x time interaction, F = 1.5, p = 0.22; effect of exposure, F 0.02, p = 0.86). **i)** Representing firing of L2/3 SOM neurons in response to increasing depolarizing current (100, 200, 400 pA current injections) 1 day after sham (grey) and noise (orange) exposure. **j)** Average firing frequency of SOM neurons as a function of injected current amplitude 1 day after sham (grey) and noise (orange) exposure. **k)** Average firing frequency as a function of injected current amplitude 10 days after sham (grey) and noise (orange) exposure. **l)** Temporal pattern of action potential generation in SOM neurons after sham (grey) and noise (orange) **m)** Average adaptation ratio (f9/f2, see panel l for traces) of SOM neurons firing rate after noise- or sham-exposure. Filled circles represent the adaptation ratio of individual SOM neurons. (Mixed model ANOVA; exposure x time interaction, F = 3.04, p = 0.08; effect of exposure, F = 0.42, p = 0.51).

We next investigated whether changes in the synaptic inputs to SOM neurons could contribute to the suppression of SOM neurons after NIHL. Despite our three-population model correctly predicting that the suppression of SOM neurons can lead to the recovery of PNs, it cannot capture such a synaptic mechanism in its current form: SOM neurons lack significant recurrent inhibition from themselves and PV neurons^33,34,66^ **(Fig. 3a)**. However, VIP neurons, which were not included in our initial model, are strongly embedded in the AC recurrent network. They have substantial incoming connections from PNs, PV, and SOM neurons, and considerable outgoing connections into SOM neurons^66,67^. Most notably, the strong mutual inhibition between SOM and VIP neurons (**Fig. 7a**; highlighted) potentially drives a competitive dynamic between these two IN subpopulations, where tipping the activity in favor of one subpopulation could lead to a dramatic suppression of the other subpopulation^68^. We, therefore, extend our computational model to include these VIP INs to investigate whether they could contribute to the observed SOM suppression.

**Figure 7.**
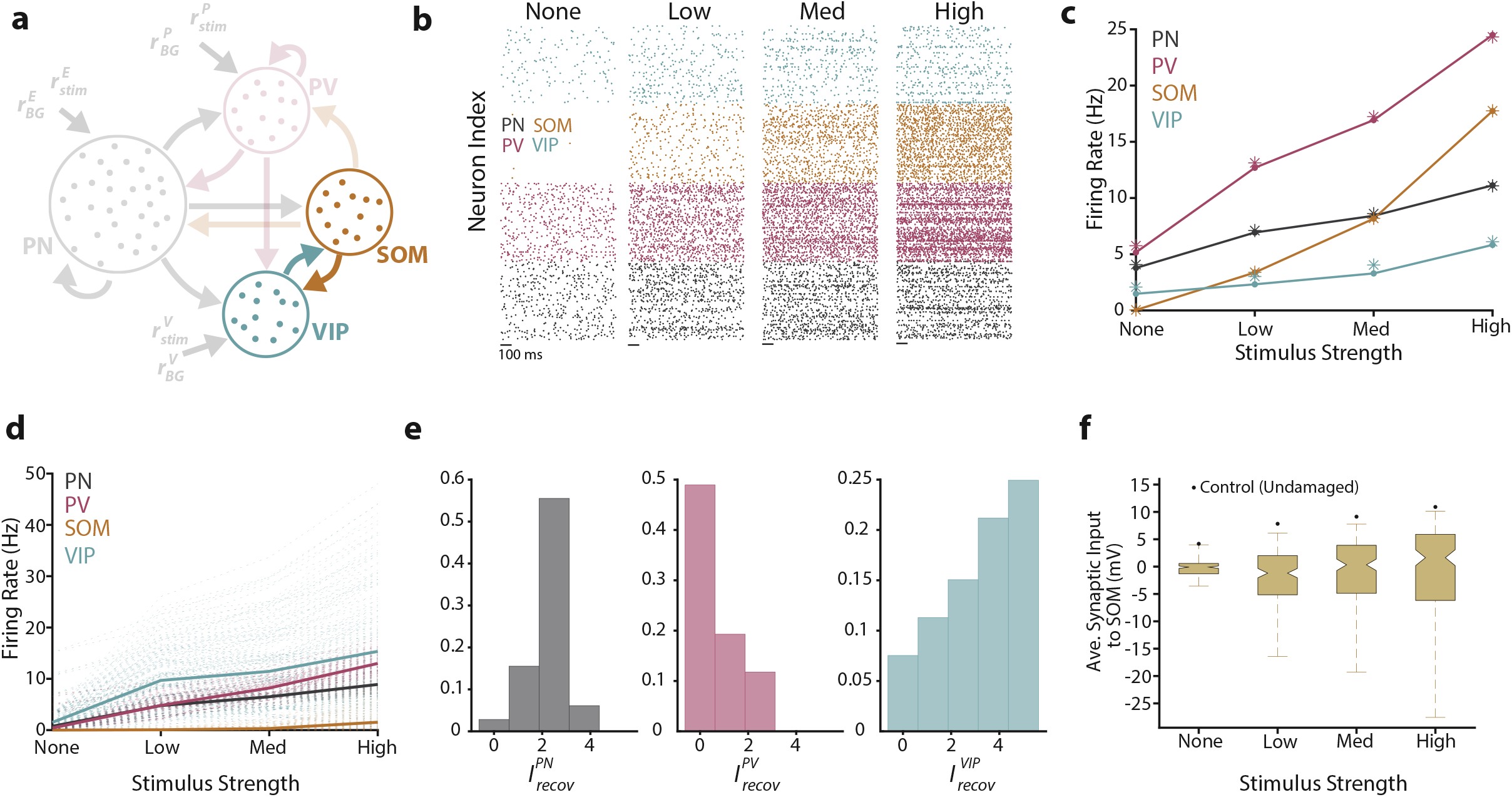
Four-population model demonstrates that VIP can lead to the threshold and gain recovery via the suppression of SOM cells. **(a)** Schematic of the four-population model, with the mutual inhibition between VIP and SOM neurons highlighted. **(b)** Raster plots showing the spiking activity of a subset of neurons at four stimulus levels for the PN (black), PV (magenta), SOM (orange), and VIP (cyan) neuron populations. **(c)** Firing rate for the spiking model (dot-line) and mean-field theory (asterisks). **(d)** Firing rate for the four populations. Translucent lines correspond to distinct parameter sets, while bolded lines are the average firing rates across all viable parameter sets. **(e)** Histograms of the recovery currents were found in the viable parameter sets for the PN, PV, and VIP neuron populations. SOM neurons did not receive a direct recovery current in this parameter search. **(f)** Box plots showing the range of average synaptic input to SOM neurons for the viable parameter sets, along with the value for the default (sham) case (black dot). All parameter values can be found in Tables 1 and 2.

For the control (pre-damaged) state, we found that the four-population model exhibited similar spiking behavior as in the three-population model **(Fig. 7b)** and that the mean-field theory was readily extendable to accurately capture the underlying steady-state firing rates **(Fig. 7c)**. With this baseline behavior in hand, we next performed a similar parameter sweep as before (see Methods for additional details). We found that the firing rates corresponding to the viable parameter sets (i.e., parameter values that yielded a low threshold, high gain, and stable dynamics for the PN population) for the PN, PV, and SOM subpopulations in the extended four-population model matched those found in our simplified three-population model (**Fig. 3e**). Specifically, the population-averaged firing rates of the PNs and PV neurons showed a low threshold and high gain, while the SOM neurons were largely suppressed (**Fig. 7d**). The difference here was that the inhibition of SOM neurons was brought on solely by VIP neurons and not by a hyperpolarizing recovery current (as was the case in the three-population model). In line with this observation, VIP neurons exhibited an increase in firing rates after damage compared to control, while also showing similar characteristics as the PNs and PV neurons, namely a low threshold and high gain (**Fig. 7d**). After examining the recovery currents responsible for these results, we observed that PN, PV, and VIP neurons were all subject to significant depolarizing currents, with VIP neurons receiving the strongest level of depolarization **(Fig. 7e)**. This result combined with the strong VIP to SOM neuron connection suggests that, during recovery after trauma, SOM neurons are more inhibited compared to the control state.

To test this directly, we measured the average synaptic input to the SOM neurons for all viable parameter sets (**Fig. 7f**, see Methods for additional details). We found that the average synaptic input was less after trauma when compared to control across all viable parameter sets and stimulus values. Further, for a majority (51.47%) of these tested conditions, SOM neurons received a net inhibitory input. In total, these modeling results from the four-population model provide a clear, testable hypothesis: VIP neurons show a strong recovery post NIHL.

To test this hypothesis experimentally, we first used *in vivo* wide-field transcranial imaging of populations of VIP neurons (**Fig. 8a-f**). We found that the response threshold of A1 VIP neurons was significantly lower than the response threshold of PNs, PV, and SOM neurons 1 day after NIHL and showed full recovery by 10 days after NIHL (**Fig. 8d**, cyan). Further, VIP neuron response amplitudes surpassed their pre-noise-exposure amplitudes 10 days after NIHL (**Fig. 8e**, cyan), and the gain was also increased throughout the 10 days after NIHL (**Fig. 8f**). We did not observe a change in the response threshold and gain of VIP neurons in sham-exposed mice (**supplement Fig. 8ab)**. Consistent with our transcranial results, longitudinal 2P imaging of individual A1 L2/3 VIP neurons, also revealed recovered (low) response thresholds, robust and even enhanced response amplitudes, and increased gain (**Fig. 8g-p**, 70 neurons from 8 mice). Also, the majority of VIP neurons showed increased gain after NIHL (**Fig. 8p insets and supplement Fig. 8g**; day1: 36/66, day3: 43/70, and day70: 47/70). On the other hand, we observed slightly reduced gain and no change in the response threshold and amplitude of A1 L2/3 VIP neurons in sham-exposed mice (60 neurons from 6 mice, **Fig. 8k, n and supplement Fig. 8c-g**). Taken together, our results support a strong recovery of the VIP neurons activity after noise trauma, even surpassing the control activity. Because VIP neurons inhibit SOM neurons^33^ (**Fig. 8b**), these results support the following circuit mechanism for cortical recovery after peripheral damage: robust VIP activity enables SOM neuron suppression, which in turn leads to high PN and PV neuron gain.

**Figure 8.**
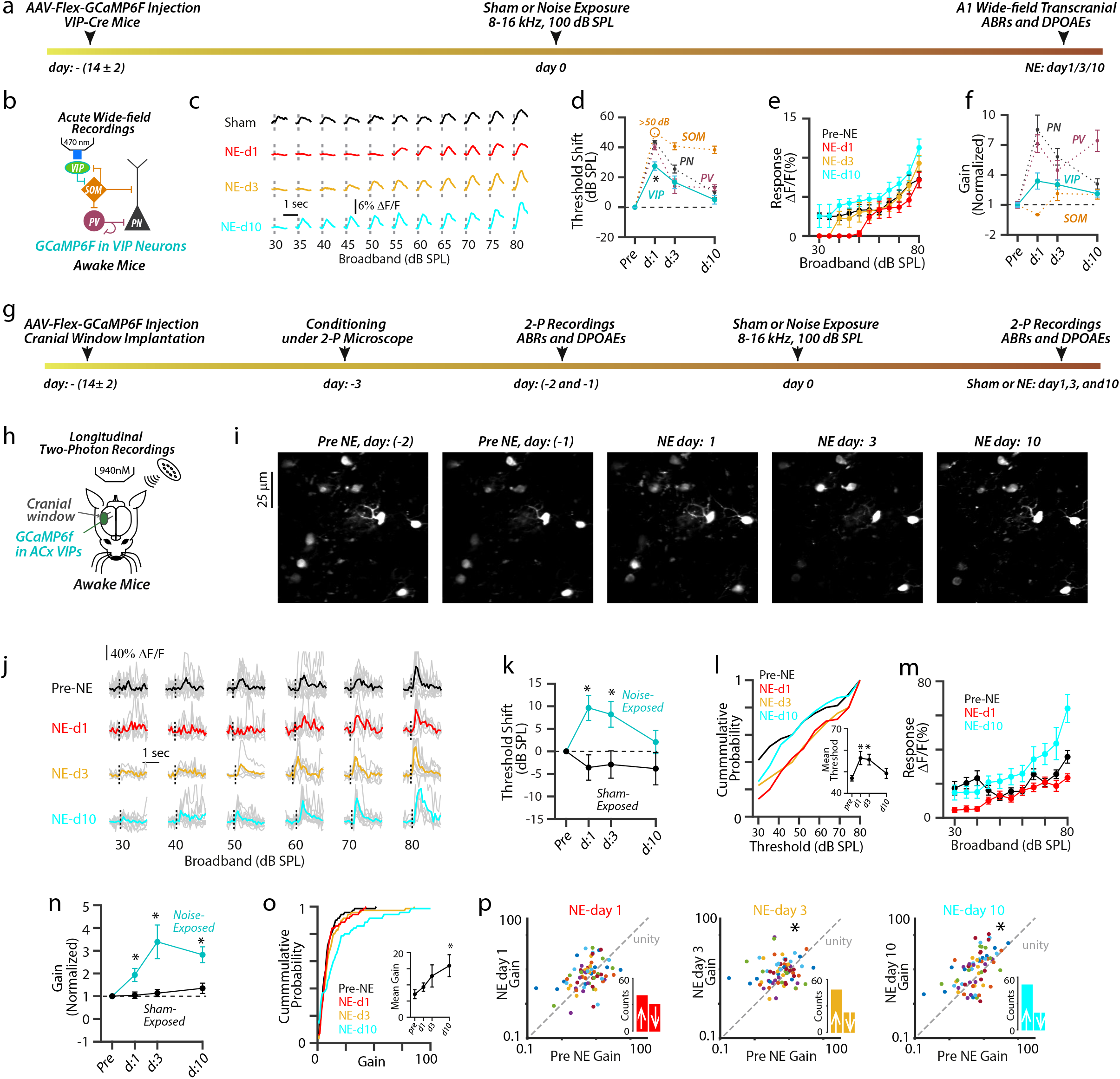
Robust sound-evoked activity (recovery) of A1 VIP neurons after NIHL. **(a)** Timetable of wide-field (WF) imaging experimental design for A1 VIP neurons. **(b)** Schematic of the cortical circuit and experimental setup illustrating transcranial imaging of A1 VIP neurons using GCaMP6f in a head-fixed awake mouse. **(c)** Representative transcranial fluorescence responses of A1 VIP neurons to broadband sounds from sham- and noise-exposed mice. **(d)** Average change in response thresholds of A1 VIP neurons (cyan) before and at 1, 3, and 10 days after noise exposure. (Noise: 4-6 mice vs. sham: 3 mice, mixed model ANOVA; exposure x time interaction, F = 8.8, p = 3.1 × 10^−4^; effect of exposure, F = 34.6, p = 3.3 × 10^−6^; *, p < 0.05, compared to pre-noise-exposed, Holm-Bonferroni’s post hoc). Average change in PN (grey), PV (red), and SOM (orange) threshold reproduced from **Figure 5**. (PV vs. PN vs. SOM vs. VIP: mixed model ANOVA; cell-type x time interaction, F = 9.3, p = 1.0 × 10^−8^; effect of time, F = 182.5, p < 10^−10^; *, p < 0.05, compared to PNs, PV, and SOM neurons at NEday1, Holm-Bonferroni’s post hoc). **(e)** Average sound-evoked responses of A1 VIP neurons to broadband sounds from noise-exposed mice. (n = 4-6, 2-way ANOVA; time x sound level interaction, F = 1.02, p = 0.44; effect of time, F = 29.66, p < 1.1 × 10^−10^. Compared to pre-noise-exposed: NE-day1, p = 5.2 × 10^−7^; NE-day3, p = 0.86, and NE-day10, p = 2.6 × 10^−4^, Holm-Bonferroni’s post hoc). **(f)** Average response gain of A1 VIP neurons (cyan), normalized to pre-noise-exposed gain, at 1, 3, and 10 days after noise exposure. (Noise: 4-6 mice vs. sham: 3 mice, mixed model ANOVA; exposure x time interaction, F = 3.1, p = 0.04; effect of exposure, F = 13.51, p = 0.007). Normalized PN (grey), PV (magenta), and SOM (orange) neuron gain reproduced from **Figure 5. (g)** Timetable of longitudinal 2-photon imaging experimental design for A1 L2/3 VIP neurons. **(h)** Schematic of experimental setup illustrating longitudinal 2-photon imaging of A1 L2/3 VIP neurons. **(i)** Z-stack images of a population of A1 L2/3 VIP neurons tracked before and after NIHL. **(j)** Representative sound-evoked responses from A1 L2/3 VIP neurons before and after NIHL. **(k)** Average change in response threshold of A1 L2/3 individual VIP neurons from noise (cyan) and sham (grey) exposed mice. (Noise-exposed: 70 neurons from 8 mice, sham-exposed: 60 neurons from 6 mice, noise vs. sham: 2-way ANOVA; exposure x time interaction, F = 3.8, p = 0.01; effect of exposure, F = 9.2, p = 0.002; *, p < 0.05, compared to pre-noise-exposed, Holm-Bonferroni’s post hoc). **(l)** Cumulative response threshold of A1 L2/3 VIP neurons, before and after NIHL. Inset: Average mean threshold of VIP neurons per mouse (Repeated measure one-way ANOVA; F = 8.3, p = 0.001. *, p < .05, compared to pre-noise-exposed, Holm-Bonferroni’s post hoc). **(m)** Average sound-evoked responses of A1 L2/3 individual VIP neurons to broadband sounds from noise-exposed mice. (2-way ANOVA; time x sound level interaction, F = 2.6, p = 1.1 × 10^−9^; effect of time, F = 39.9, p < 1.1 × 10^−10^; Compared to pre-noise-exposed: NE-day1, p = 5.1 × 10^−9^; NE-day3, p = 0.004 and NE-day10, p = 1.1 × 10^−5^, Holm-Bonferroni’s post hoc). **(n)** Average gain of A1 L2/3 VIP neurons normalized to pre-exposed gain from noise (cyan) and sham (grey) mice. (Noise vs. sham: 2-way ANOVA; exposure x time interaction, F = 4.7, p = 0.002; effect of exposure, F = 13.7, p = 3.1 × 10^−4^*, p < 0.05, compared to pre-noise-exposed, Holm-Bonferroni’s post hoc). **(o)** Cumulative gain of A1 L2/3 VIP neurons, before and after NIHL. Inset: Average mean gain of PNs per mouse (Friedman test; friedman statistic = 9.4, p = 0.02. *, p < .05, compared to pre-noise-exposed, Holm-Bonferroni’s post hoc). **(p)** Scatter plots of the gain of individual A1 L2/3 VIP neurons before and after NIHL. Dotted line represents unity. Insets: Bar graphs representing the number of neurons showing increased gain (↑ above unity) and reduced gain (↓ below unity) after NIHL. PreNE vs. NEday1: p = 0.4, PreNE vs. NEday3: p = 0.04, and PreNE vs. NEday10: p = 3.9 × 10^−4^; permutation test.

## Discussion

### Division-of-labor between the cortical IN subtypes

Extensive evidence supports divergence, complementarity, and division-of-labor between the cortical IN subtypes in terms of their tuning properties^69-72^, and their role in contextual and adaptive cortical sound processing^40,43,73,74^. However, despite the established role of reduced GABAergic signaling in A1 plasticity after cochlear damage^11,22,26,27,75,76^, the roles of different IN subtypes in cortical recovery remained unknown. We found that while auditory nerve input to the brainstem is significantly reduced after cochlear damage, sound-evoked cortical activity is maintained or even enhanced. Importantly we revealed a strategic, cell-type-specific, and time-dependent plasticity scheme that restores cortical responses. Namely, after noise trauma, we found enhanced sound-evoked activity in PN **(Fig. 2)**, PV **(Fig. 4)**, and VIP neurons **(Fig. 8)**, but reduced sound-evoked activity in SOM neurons **(Fig. 5)**. Based on the known sequentially organized inhibitory cortical network^33^, where VIP neurons inhibit SOM neurons, SOM neurons inhibit PV neurons and PNs, and PV neurons inhibit PV neurons and PNs, we propose that the underlying SOM → PV → PN and SOM → PN circuits support a cell-specific plasticity mechanism in which, robust PV activity provides network stability by balancing PN activity; and vastly decreased SOM IN activity allows for increased PV IN and PN gain, which supports stability and high gain. The VIP → SOM → PN disinhibitory pathway completes the task, whereby robust VIP IN activity enables reduced SOM IN activity. These results highlight a novel strategic, cooperative, and cell type-specific plasticity program that restores cortical sound processing after cochlear damage and provides novel cellular targets that may also aid in the development of pharmacotherapeutic or rehabilitative treatment options for impaired hearing after NIHL.

Several key cortical circuit features are consistent with our proposed hypothesis, regarding the roles of PV neurons as stabilizers and SOM neurons as modulators of A1 plasticity after NIHL. PN and PV neurons are embedded into very similar synaptic environments. Both receive the excitatory drive from upstream areas^33,34^, and both receive strong recurrent excitation, as well as PV- and SOM-mediated inhibition^66^. This symmetry places PV neurons in a strategic position for monitoring and stabilizing PNs activity^53^. On the other hand, SOM neurons are in a better position to modulate the cortical inhibition and excitation^53^, such that a higher gain state of PNs can be achieved without compromising the stability of the network. For example, the increased gain of PNs via reduced activity of SOM neurons would lead to the increased firing of PV neurons because of the strong excitatory feedback from PNs to PV neurons. In turn, these hyperactive PV neurons would then stabilize the recurrently activated PNs. Further, consistent with our model predictions **(Fig. 3e)**, suppression of SOM neurons enhances cortical plasticity without compromising the stability of the network^52,77,78^, whereas suppression of PV neurons can result in uncontrolled network activity, evidenced by unstable ictal-like events in most^52^, but not all cases^79^. These observations support the notion that PV neurons act as the stabilizers whereas SOM neurons act as the modulators of A1 plasticity.

Similarly, our proposed role of VIP neurons as the enablers of A1 plasticity after NIHL is consistent with previous reports showing that VIP neurons enable cortical plasticity across the sensory cortices^68,74,77,78,80-82^. In the visual cortex, synaptic transmission from VIP to SOM neurons is necessary and sufficient for the increased cortical responses in PNs after monocular deprivation in adult mice^77^. In the somatosensory cortex, increased activity of VIP neurons facilitates the increased activity of PNs in a mouse model of neuropathic pain^83^. Here, we found the enhanced activity of A1 L2/3 VIP neurons after NIHL **(Fig. 8)**, suggesting that VIP neurons enable the enhanced activity of PNs **(Fig. 2)** via the disinhibitory pathway VIP → SOM → PN. Together, our results provide the first comprehensive, and precise cell-type- and circuit-mechanism for how cortex rebuilds itself after peripheral damage.

### Maladaptive aspects of cortical plasticity

Compensatory plasticity in the A1 after peripheral damage supports the recovery of the perceptual sound-detection threshold but does not support sound processing encoded by precise spike timing, such as modulated noise or speech and restricts hearing in a noisy environment^10,11,15,84,85^. Interestingly, A1 SOM neurons, which are critically important for sound processing encoded by precise spike timing of neuronal firing^43,44,86,87^, showed reduced sound-evoked activity after NIHL **(Fig. 5)**. Based on these results, we propose that the reduced activity of A1 SOM neurons after cochlear damage may contribute to the hearing problems peripheral damage, such as difficulty in understanding speech and trouble hearing in noisy environments.

Another maladaptive aspect of increased AC gain after peripheral damage is the development of tinnitus, the perception of phantom sounds^15^, and hyperacusis, the painful sensitivity to everyday sounds^16^. Both these disorders have many similarities with neuropathic pain and phantom limb syndrome^88^. These neurological disorders are developed after damage to the peripheral organs and manifest increased activity of PNs in the respective sensory cortices, suggesting a common underlying cortical circuit mechanism. In the allodynia mouse model of neuropathic pain^83^, where sensory touch that does not normally provoke pain becomes painful in the spared nerve injury model, SOM activity was drastically reduced in the somatosensory cortex. This is consistent with the notion that reduced activity of SOM neurons disinhibits the PNs and leads to increased activity of PNs. Interestingly, selective activation of SOM neurons in the somatosensory cortex after nerve injury was sufficient to prevent the increased activity of PNs and to mitigate the development of neuropathic pain^83^. Together, these results suggest that the reduced activity of cortical SOM neurons after peripheral organ damage may be a common mechanism across sensory cortices that permits for the increased gain of PNs. Moreover, these results suggest that modulation of A1 SOM neuron activity after noise trauma could be a potential target for mitigating noise-induced tinnitus and hyperacusis.

### PV neurons and cortical plasticity after peripheral trauma

Our results suggest that the increased sound-evoked activity of PV neurons 10 days after NIHL **(Fig. 4d and k, cyan)** may stabilize the increased activity of PNs **(Fig. 2)**. However, 1 day after NIHL we observed reduced sound-evoked PV neuron activity **(Fig. 4d and k, red)**. Since, PV neurons initiate cortical plasticity in juvenile and adult brain^85^, an initial reduction in the activity of PV neurons activity after NIHL may initiate cortical plasticity. Consistent with this notion, a rapid drop in PV-mediated inhibition of PNs as early as 1 day after cochlear denervation precedes the recovery of cortical sound processing^10^. In the visual cortex, PV neurons show reduced firing rates 1 day after monocular deprivation^80^, and in the somatosensory cortex PV neurons also show reduction in their intrinsic excitability 1 day after whisker plugging^89^. These results suggest that a rapid reduction in PV-mediated inhibition of PNs may be a common feature of sensory cortices plasticity that plays a critical role in initiating cortical recovery after sensory organ damage.

A recent study^11^ reported that the sound-evoked activity of PV neurons was reduced after cochlear denervation and remained reduced for two weeks. However, unlike our model of noise-induced cochlear damage, cochlear denervation was induced with bilateral cochlear application of ouabain, which eliminates ∼95% of the type 1 SGNs. Since the type and the severity of peripheral organ damage may result in heterogeneous cortical plasticity^7,10,16,22,29,90-96^, noise- and ouabain-induced damage to the cochlea may trigger different trajectories of plasticity in A1 cell-types. Another explanation for the observed differences could arise from differences in the experimental design, such as the sound-stimuli used (broadband vs. 12 kHz pure tones^11^) and the number of PV neurons tracked (82 vs 29^11^). Overall, the observed differences point to the need for further rigorous investigations on the role of distinct INs in A1 plasticity in different types and degrees of hearing loss, including unilateral vs. bilateral land noise vs. ototoxic compounds induced hearing loss. Nonetheless, our results provide a comprehensive model of cortical rehabilitation after noise trauma whereby the precise and well-timed division-of-labor and cooperativity among cortical interneurons secure high gain and stability.

### Model assumptions and limitations

In this work, we leveraged a computational model to assist with exploring cortical mechanisms responsible for the recovery of PNs following NIHL. We modeled recovery at the neuronal level as either depolarization or hyperpolarization of the resting membrane potential. After peripheral injury such as NIHL, it is plausible to assume that homeostatic mechanisms could activate such mechanisms, either intrinsically or synaptically, in an attempt to return neuronal populations to previous baseline levels. However, of the two, it is perhaps less likely that mechanics would lead to a hyperpolarization, as was predicted for SOM neurons since this would lead to further population suppression. Despite this shortcoming, the experimental data confirmed our model’s prediction. It also suggested that this current arose from synaptic pathways, which pointed to a possible model update: include VIP neurons within the model AC circuit. This extension of our computational model allowed it to capture the experimental results with the use of only depolarization of neuronal membrane post-injury. However, it remains an open question as to the exact source of such currents.

The computational model assumes that after NIHL, the network must balance the mechanics of recovery and stabilization. This assumption, namely that cortical inhibition is needed to prevent pathologic activity, places the model in the inhibition-stabilized network (ISN) regime^46,51,53,62^. While we have already mentioned the evidence pointing to PV neurons as the best IN subtype to play the role of the stabilizer, after suffering NIHL, synaptic plasticity and other mechanisms may divert this role to an alternative IN or shift the entire circuit out of the ISN regime. If this were the case, the region of viable parameter sets for the computational model would grow and, as a result, would suggest alternative recovery pathways (e.g., hyperpolarizing PV neurons). Yet, the absence of such pathways in our experimental results implies that the dynamics observed in the ISN regime constrain the mechanisms utilized in recovery. Whether the cortex lies in the ISN regime post-NIHL or remains wired to resist instability despite no longer requiring it remains an open question.

In conclusion, our results create a new framework for understanding the cellular and circuit mechanisms underlying AC plasticity after peripheral trauma and hold the promise to advance understanding of the cortical mechanisms underlying disorders associated with maladaptive cortical plasticity after peripheral damage, such as tinnitus^14,15^, hyperacusis^16,59^, visual hallucinations^97^, and phantom limb pain^13,98^.

## Methods

### Animals

All procedures were approved by the Institutional Animal Care and Use Committee at the University of Pittsburgh. For experiments shown in Figure 1, male and female C57/B6 mice and PV-Cre, SOM-Cre mice, and VIP-Cre mice with C57/B6 mice backgrounds (The Jackson Laboratory) were used. For experiments shown in Figures 2 and 4, male and female PV-Cre mice were used. For experiments shown in Figure 5, male and female SOM-Cre were used. For experiments shown in Figure 6, male and female SOM-GFP (GIN) mice with C57/B6 mice backgrounds were used. For experiments shown in Figure 8, male and female VIP-Cre mice were used.

### Speaker Calibration

Acoustic sound stimuli used in the study were calibrated with pre-amp attached microphones (1/8 inch 4138-A-015 and 1/4 inch 4954-B, Brüel and Kjær) and a reference 1 kHz, 94 dB SPL certified speaker (Type 4231, Bruel & Kjaer). More specifically, we placed the microphone at the same position as the mouse ear and delivered the pure tones and broadband stimuli at a specific voltage input and recorded output voltage using the pre-amp microphone. Then, we determined the voltage input needed to generate the desired dB SPL output using the 1 kHz 94 dB SPL speaker as the reference voltage.

### Behavioral Training and Testing

Behavioral training and testing were performed in a shuttle-cage (14” W x 7” D x 12” H) bisected into two virtual zones^99^. The shuttle-cage was placed within a sound- and light-attenuation chamber and was equipped with 8 poled shocking floor, calibrated speaker, and mouse position sensors (Coulbourn Instruments). Sound-stimuli were generated using a programable tone/noise generator (A12-33, Coulbourn Instruments) and delivered via calibrated multi-field speaker (MF1, Coulbourn Instruments) hung in the middle of the shuttle cage to provide a homogenous sound field. Foot-shock signals were generated using a programable animal shocker (H13-17A, Coulbourn Instruments). Presentation of sound-stimuli, foot-shock signals and mouse position detection were performed using scripts programmed in GRAPHIC STATE 4 (Coulbourn Instruments).

On the day of training and testing, mice were given at least 5 min to acclimate to the shuttle-cage before beginning the training or testing session. Six to seven-week-old C57B6 mice were initially trained to cross from one side of the shuttle-box to another side to terminate a 200 μA foot-shock. Foot-shock was terminated upon crossing to the other side of the shuttle-box or 10 seconds, whichever occurred first. Seven to eight blocks of 10-foot-shock trials were performed during a training session for two days. Next, mice were trained to cross from one side of the shuttle-box to the opposite side following a sound-stimuli (50 ms noise-burst, 6-sec long train with a repetition rate of 2.5 Hz at 70 SPL, with a randomized intertrial interval of 30-40 sec) to avoid a mild foot-shock (200 - 400 μA)A successful crossing during the noise-bursts trial was called Hit, whereas crossing during a random 6-sec long silent window was called False-Alarm (FA). Seven to eight blocks of 10 sound-stimuli trials were performed during a training session every day for 4-6 days. Mice were trained every day until their behavioral performance d’ exceeded the value of 1.5 [d’ = z-score(Hit rate) – z-score(FA rate)]. During the behavioral testing sessions, we presented noise-bursts at various sound intensity levels (20-80 dB SPL) in random order. Each sound level was presented at least 10 times with a randomized intertrial interval of 30-40 sec. We measured the Hit and FA rates at individual sound-level and calculated the sound-detection rate (Hit% - FA%). To quantify the sound-detection threshold, we plotted the sound-detection rate against the sound-levels, and the sound-level with a 50% sound-detection rate was defined as the sound-detection threshold.

### Noise Exposure

Unanesthetized and unrestrained mice were placed within a 5×4-inch acoustically transparent box, and bilaterally exposed to an octave band (8-16 kHz) noise at 100 dB SPL for 2 hours noise from a calibrated speaker. For sham-exposed mice, unanesthetized and unrestrained mice were placed within the same box for 2 hours, but the noise was not presented.

### Auditory brainstem responses

Auditory brainstem response (ABR) thresholds and ABR wave 1 amplitude were measured with subdermal electrodes in mice under isoflurane anesthesia at a stable temperature (∼37° C) using the RZ6 processor (Tucker-Davis Technologies, Kumar al., 2019). We recorded ABRs after presenting broadband clicks (1 ms duration, 0 – 80 dB SPL in 10 dB steps) at a rate of 18.56 per second with a calibrated MF1 speaker (Tucker-Davis Technologies), via a probe tube inserted in the ear canal. We presented each sound 512 times and analyzed the average evoked potential after bandpass filtering the waveform between 300 and 3000 Hz. ABR threshold was defined as the lowest sound intensity that generated ABR wave I amplitudes that were 3 SDs above the baseline noise level. Baseline noise levels were measured using the ABRs obtained at 0 dB SPL sound intensity. ABR wave 1 amplitude was measured from peak to trough levels.

### Distortion product otoacoustic emissions

Mice were anesthetized using isoflurane (3% Induction/ 1.5% Maintenance, in oxygen) and kept at a stable temperature using a heating pad (∼37° C). Measurements for DPOAE thresholds were taken with the RZ6 processor and BioSigRX software (Tucker-Davis Technologies). Tone pairs were presented with an f1 and f2 primary ratio of 1.2 at center frequencies. The f1 and f2 primaries were presented using 2 separate MF1 speakers (Tucker-Davis Technologies) that each presented a frequency into the outer ear canal, by using tubing that came together within an acoustic probe to limit artificial distortion. The presentation of these tones into the cochlea results in a distortion product, which is generated by the outer hair cells and recorded by a sensitive microphone. Recordings were taken at 8, 12,16, 20, and 24 kHz in ascending order from 0-80 dB. Each test frequency and intensity were averaged over one hundred sweeps. DPOAE threshold was determined as the lowest intensity that was able to generate a distortion product (2f1-f2) with an amplitude that was at least three standard deviations above the noise floor.

### Adeno-associated virus injections for in vivo imaging

Male or female PV-Cre mice, SOM-Cre mice, and VIP-Cre mice (The Jackson Laboratory) were injected with AAV9.CaMKII.GCaMP6f. WPRE.SV40 and AAV9.CAG.Flex.GCaMP6f.WPRE.SV40 into the right auditory cortex as described previously^65,100,101^. Briefly, mice were anesthetized with isoflurane, and a craniotomy (∼0.4 mm diameter) was made over the temporal cortex (∼4 mm lateral to lambda). With a micromanipulator (Kopf), a glass micropipette containing AAVs was inserted into the cortex 0.5–0.7 mm past the surface of the dura and ∼500 nL of each viral vector was injected over 5 min. Next, the scalp of the mouse was closed with cyanoacrylate adhesive. Mice were given carprofen 5 mg/kg (Henry Schein Animal Health) to reduce the pain associated with the surgery and monitored for signs of postoperative stress and pain.

### Animal preparation for acute in *vivo* wide-field imaging

Twelve to 16 days after AAV injections, mice were prepared for *in vivo* calcium imaging^65,100,101^. Mice were anesthetized with inhaled isoflurane (induction, 3% in oxygen; maintenance, 1.5% in oxygen) and positioned into a custom-made head holder. Core body temperature was maintained at ∼37**°**C with a heating pad, and eyes were protected with ophthalmic ointment. Lidocaine (1%) was injected under the scalp, and an incision (∼1.5 cm long) was made into the skin over the right temporal cortex. The head of the mouse was rotated ∼45° in the coronal plane to align the pial surface of the right temporal cortex with the imaging plane of the upright microscope optics. The skull of the mouse was secured to the head holder using dental acrylic (Lang) and cyanoacrylate adhesive. A tube (the barrel of a 25 ml syringe or an SM1 tube from Thorlabs) was placed around the animal’s body to reduce movement. A dental acrylic reservoir was created to hold warm (37°C) ACSF over the exposed skull. The ACSF contained (in mM)130 NaCl, 3 KCl, 2.4 CaCl2, 1.3 MgCl2, 20 NaHCO3, 3 HEPES, and 10 D-glucose, pH 7.25–7.35, ∼300 mOsm. For better optical access to the auditory cortex, we injected lidocaine– epinephrine (2% lidocaine,1:100,000 w/v epinephrine) into the temporal muscle and retracted a small portion of the muscle from the skull. Mice were then positioned under the microscope objective in a sound- and light-attenuation chamber containing the microscope and a calibrated speaker (ES1, Tucker-Davis).

### *In vivo* wide-field imaging

We performed transcranial imaging to locate the primary auditory cortex (A1) and image sound-evoked activity from specific populations of A1 neurons in awake mice. We removed the isoflurane from the oxygen flowing to the animal and began imaging sound-evoked responses after 60 min of recovery from isoflurane^65,100,101^. Sounds were delivered from a free-field speaker 10 cm from the left ear of the animal (ES1 speaker, ED1 driver, Tucker-Davis Technologies), controlled by a digital-to-analog converter with an output rate of 250 kHz (USB-6229, National Instruments). We used ephus^102^ to generate sound waveforms and synchronize the sound delivery and image acquisition hardware. We presented 6 or 32 kHz, 50 dB SPL tones to the animal while illuminating the skull with a blue LED (nominal wavelength, 490 nm; M490L2, Thorlabs). We imaged the change in green GCaMP6f emission with epifluorescence optics (eGFP filter set, U-N41017, Olympus) and a 4x objective (Olympus) using a cooled CCD camera (Rolera, Q-Imaging). Images were acquired at a resolution of 174 × 130 pixels (using 4x spatial binning, each pixel covered an area of 171.1 μm2 of the image) at a frame rate of 20 Hz to locate A1 in each animal (see below, Analysis). To locate the A1, we presented low-frequency tones (5 or 6 kHz, 40–60 dB SPL) and imaged the sound-evoked changes in transcranial GCaMP6s fluorescence. Due to the mirror-like reversal of tonotopic gradients between A1 and the anterior auditory field (AAF)^64,103^, these sounds activated two discrete regions of the auditory cortex corresponding to the low-frequency regions of A1 and the AAF **(supplement Fig. 2a)**. To extract change sound-evoked change in fluorescence (ΔF/F), we normalized the sound-evoked change in fluorescence after the sound presentation (ΔF) to the baseline fluorescence (F), where F is the average fluorescence of 1 s preceding the sound onset (for each pixel in the movie). We applied a two-dimensional, low-pass Butterworth filter to each frame of the ΔF/F movie and created an image consisting of a temporal average of 10 consecutive frames (0.5 s) beginning at the end of the sound stimulus. After localizing A1, we presented broadband sounds (6-64 kHz, 100 ms long) at 30-80 db SPL in 5 db SPL steps from a calibrated speaker (ES1, TDT) and imaged the sound-evoked changes in transcranial GCaMP6f fluorescence signals (ΔF/F%). Each sound was presented 8-10 times in pseudo-random order.

### *In vivo* wide-field imaging analysis

A region of interest (ROI, 150–200 mm x 150–200 mm) over A1 was then used to quantify the sound-evoked responses to broadband sounds (6-64 kHz, 100 ms long) sounds. We averaged the fluorescent intensity from all pixels in the ROI for each frame and normalized the ΔF to the F of the ROI to yield ΔF/F responses. ΔF/F responses from 8 to 10 presentations of the same sound level were averaged. Response amplitude was the peak (50 msec window) of the transcranial response that occurred within one second of the sound onset. Response threshold was defined as sound-level which elicit a significant increase in fluorescent signals (2 standard deviations above baseline fluorescence F). The response gain was defined as the slope of response amplitudes against the sound levels and calculated as the average change in the fluorescence signals (ΔF/F%) per 5 dB SPL step starting from response threshold^10,11^.

### Longitudinal *in vivo* two-photon imaging

After AAV injections into the right AC as described above, we implanted a 3 mm wide cranial glass window over the AC following a published protocol^80,104^. A metal head-plate was also affixed to the mice’s heads using dental cement to hold them under the 2P microscope. Twelve to 16 days after the surgery, mice were first conditioned/habituated under the 2P microscope. Mice were head-fixed under a 2P microscope with the head-plate and allowed to acclimate to the rig set up for 30-40 minutes while we passively played broadband and pure-tone sounds in the background. The next day, after locating A1 using wide-field imaging as described above, we performed two-photon imaging of A1 L2/3 neurons (175-225 μm below the pial surface) in awake mice. Mode-locked infrared laser light (940 nm, intensity at the back focal plane of the objective, MaiTai HP, Newport, Santa Clara, CA) was delivered through a galvanometer-based scanning two-photon microscope (Scientifica, Uckfield, UK) controlled with scanimage 3.8^105^, using a 40x, 0.8 NA objective (Olympus) with motorized stage and focus controls. We imaged green and red fluorescence simultaneously with two photomultiplier tubes behind red and green emission filters (FF01-593/40, FF03-525/50, Semrock) using a dichroic splitter (Di02-R561, Semrock) at a frame rate of 5 Hz over an area of 145 × 145 μm and at a resolution of 256 × 256 pixels. We imaged PNs, PV, SOM, and VIP neurons in L2/3 at a depth of ∼200 μm from pia. Next, we presented trains of broadband sounds at interstimuli intervals of 5 s (6-64 kHz, 100 ms long) at 30-80 db SPL in 5 db SPL steps in pseudo-random order and imaged the sound-evoked changes in GCaMP6f fluorescence signals (ΔF/F%). The whole two-photon imaging session lasted 20-30 mins long and upon completion, the mice were returned to their cage. To use each neuron as its own control, we manually tracked the same neurons for 10 days after noise- or sham-exposure and imaged sound-evoked changes in GCaMP6f fluorescence signals to sound-trains. Mice were habituated under the 2-photon objective for 20-30 minutes a day before the pre-exposure recording sessions. Pre-exposure sessions lasted two days and average responses of individual neurons from both days were used as pre-exposure responses.

### Two-Photon Analysis

Images were analyzed post hoc using a custom program, and open-source routines, written using Python and MATLAB as described previously^106^. Before extracting ΔF/F, we used the NoRMCorre software to correct motion artifacts from individual tiff movies^107^. Next, using FISSA: A neuropil decontamination toolbox for calcium imaging signals^108^, we selected ROIs around the soma of each L2/3 neuron from the temporal average of all tiff movies from a single recording session. Fluorescence values were extracted from each ROI for each frame, and the mean for each cell was computed. FISSA, gave us two vectors of fluorescence values for the somatic and the neuropil. We weighted the neuropil vector by 0.8 as described previously^106,109,110^. The weighted neuropil vector was subtracted from the somatic vector to produce a corrected vector of fluorescence values. These FISSA corrected fluorescence (F) value from each sound trial were then converted to ΔF/F values by using baseline fluorescence measured 1 sec before each sound onset. We then averaged the ΔF/F values from each sound-trail (5-8 trials) to get mean ΔF/F from each neuron. To identify the sound-responsive neurons, we used a tone sensitivity index, d-prime (d’), from preexposure sessions as described previously^103,106,111^. Briefly, we presented trains of pure tones at interstimuli intervals of 3 s in pseudo-random order that spanned in the range of 4–40 kHz frequencies (500-ms long) in 0.20-octave increments at 30-80 dB SPL in 10 dB SPL steps. For each neuron, we calculated the average response amplitude from responses at and immediately adjacent to the frequency/level combination eliciting the maximum response (average of 5 values if the maximum response is observed at dB < 80, 4 values if the maximum response is observed at 80 dB). We then averaged the same number of values selected at random frequency/level locations of the frequency response area (FRA). We took the difference between these averages and iterated this process 1000 times. The tone sensitivity index, d-prime (d’), was calculated as the average of the iterated differences, and the neurons with d’ ≥ 0 only were analyzed further. We then used these sound-responsive cells to assess sound-evoked activities, such as response threshold, amplitude, and gain. The sound-evoked responses were measured for 1 s after the sound onset and were defined as significant responses if the sound-evoked changes in ΔF/F were larger than the mean+2 standard deviations (SDs) of the baseline fluorescence measured before the sound onset. Peak fluorescence signals during the 1-s period after the sound presentation were quantified as the sound-evoked response amplitude. The response gain was defined as the slope of response amplitudes against the sound levels and calculated as the average change in the fluorescence signals (ΔF/F%) per 5 dB SPL step starting from response threshold^10,11^.

### Brain slice *ex vivo* electrophysiology

We recorded intrinsic properties of AC SOM neurons as described previously^64,65^. Due to the lack of cytoarchitectural features, it is challenging to locate the AC in brain slices. Therefore, to localize the AC, we labeled AC corticocollicular (CCol) L5B PNs (red) projecting to the inferior colliculus, by injecting red fluorescent retrograde microspheres into the inferior colliculus of SOM-GFP mice. Briefly, P28-35 male or female, SOM-GFP (GIN) mice were injected with red fluorescent retrograde microspheres into the ipsilateral inferior colliculus (IC) (1 mm posterior to lambda and 1mmlateral, injection depth 0.75 mm). A volume of ∼0.12 μL of microspheres was pressure-injected (25 psi, 10–15 ms duration) from capillary pipettes (Drummond Scientific) with a Picospritzer (Parker–Hannifin). The localization of CCol PNs in the AC (**Fig. 6b**), along with anatomical landmarks, such as the rhinal fissure and the underlying hippocampal formation allowed us to locate the AC as described previously^63-65^. On the day of recordings, brains were rapidly removed and coronal slices (300 μm) containing the right AC were prepared in a cutting solution at 1 °C using a Vibratome (VT1200 S; Leica). The cutting solution, pH 7.4, ∼300 mOsm, contained the following (in mM): 2.5 KCl, 1.25 NaH2PO4, 25 NaHCO3, 0.5 CaCl2, 7 MgCl2, 7 glucose, 205 sucrose, 1.3 ascorbic acid, and 3 sodium pyruvate (bubbled with 95% O2/5% CO2). The slices were immediately transferred and incubated at 34 °C in a holding chamber for 40 min before recording. The holding chamber contained artificial cerebrospinal fluid (ACSF), pH 7.4, ∼300 mOsm containing the following (in mM): 125 NaCl, 2.5 KCl, 26.25 NaHCO3, 2 CaCl2, 1MgCl2, 10 glucose, 1.3 ascorbic acid, and 3 sodium pyruvate, pH 7.4, ∼300 mOsm (bubbled with 95% O2/5% CO2). Next, whole-cell recordings in voltage- and current-clamp modes were performed on slices bathed in carbogenated ACSF, which was identical to the incubating solution. For electrophysiological recordings, we used a MultiClamp-700B amplifier equipped with Digidata-1440A A/D converter and Clampex (Molecular Devices).

Data were sampled at 10 kHz and filtered at 4 kHz. To study the intrinsic properties of SOM neurons, we added the following drugs: 20 μM DNQX (AMPA receptor antagonist), 50 μM APV (NMDA receptor antagonist), and 20 μM SR 95531 Hydrobromide (Gabazine—a GABAA receptor antagonist). Pipette capacitance was compensated and series resistance for recordings was lower than 15M*Ω*. Series resistance (*R*_series_) was determined by giving a −5-mV voltage step for 50 ms in voltage-clamp mode (command potential set either at −70 mV or at 0 mV) and was monitored throughout the experiments. *R*_series_ was calculated by dividing the −5 mV voltage step by the peak current value generated immediately after the step in the command potential. Recordings were excluded from further analysis if the series resistance changed by more than 15% throughout the experiment. Input resistance (*R*_input_) was calculated by giving a −5-mV step in voltage-clamp mode (command potential set either at −70 mV or at 0 mV), which resulted in transient current responses. The difference between baseline and steady-state hyperpolarized current (*ΔI*) was used to calculate *R*_input_ using the following formula: *R*_input_= (−5 mV/*ΔI*) − *R*_series_. The average resting membrane potential (*V*_m_) was calculated by holding the neuron in voltage-follower mode (current clamp, at *I* = 0) immediately after breaking in and averaging the membrane potential over the next 20 s. In the current clamp, depolarizing current pulses (0–450 pA in 50 pA increments of 1-s duration) were used to examine each neuron’s basic suprathreshold electrophysiological properties (baseline *V*m was maintained at −70 mV). Action potential (AP) width was calculated as the full width at the half-maximum amplitude of the first resulting AP at rheobase. The AP threshold was measured in the phase plane as the membrane potential at which the depolarization slope exhibited the first abrupt change (*Δ*slope *>*10 V/s). The adaptation ratio was calculated by dividing the instantaneous frequency between the ninth and tenth AP by the instantaneous frequency between the second and third AP (f9/f2).

### Cochlear Immunohistochemistry

Mice were deeply anesthetized with isoflurane and sacrificed by decapitation within one week of behavior testing. Cochleas were extracted and perfused intralabrynthly with 4% paraformaldehyde in 0.1 M phosphate buffer. Cochleas were post-fixed for 2 hr at room temperature and decalcified in 120 mM EDTA for 2-3 days at room temperature on a rocker. Decalcified cochleas were then microdissected under a stereomicroscope. Cochlear sections were blocked in 5% normal goat serum with 0.3% Triton X-100 in phosphate-buffered saline (PBS) for 1 hr at room temperature. Sections were then incubated in primary antibodies diluted in blocking buffer overnight (18-24 hr) at room temperature. Primary antibodies used were anti-myosin VIIa (rabbit anti-MyoVIIa; Proteus Biosciences; 1:500), anti-C-terminal binding protein 2 (mouse anti-CtBP2 IgG1; BD Biosciences; 1:200), and anti-glutamate receptor 2 (mouse anti-GluR2 IgG2a; Millipore; 1:2000). Sections were then washed with PBS and incubated in Alexa Fluor-conjugated fluorescent secondary antibodies (Invitrogen; 1:500) for 2 hr at room temperature. Sections were again washed in PBS and finally mounted on microscope slides using Prolong Diamond Antifade Mountant (Invitrogen).

Cochlear sections were imaged in their entirety at low magnification to reconstruct the cochlear frequency map using an ImageJ plugin provided by Eaton Peabody Laboratories. http://www.masseyeandear.org/research/otolaryngology/investigators/laboratories/eaton-peabody-laboratories/epl-histology-resources/imagej-plugin-for-cochlear-frequency-mapping-in-whole-mounts. This preparation allows us to trace the organ of Corti in its entirety from base to apex, and the plugin superimposes the frequency map on the traced sections. Confocal z-stacks (0.25 mm step size) of the 8, 12, 16, 24, and 32 kHz regions from each cochlea were captured using a Nikon A1 microscope under a 60x oil immersion lens. Images were imported to ImageJ imaging software for quantification, where maximum projections were rendered from the z-stacks. CtBP2 and GluR2 puncta were counted to identify intact ribbon synapses. Synapses were only considered intact if CtBP2 and GluR2 puncta were juxtaposed. Orphan synapses were defined as CtBP2 puncta that lacked GluR2 puncta. Between 14-18 inner hair cells were included for synapse quantification.

### Computation Modelling, LIF network

We consider a four (*a* = *PN, PV, SOM*, and *VIP*) population network of leaky integrate-and-fire (LIF) neurons, where the membrane potential 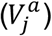 of the *j*^*th*^ neuron in population *a* is governed by the equation

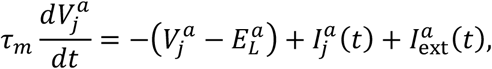

where 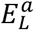 is the resting potential, *τ*_*m*_ is the membrane time constant, and 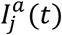 and 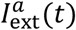 are the recurrent and external synaptic currents, respectively. When 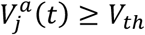, its value is reset to *V*_*r*_ and undergoes a refractory period of length *τ*_*r*_.

The recurrent synaptic currents are modeled with exponentially decaying synapses

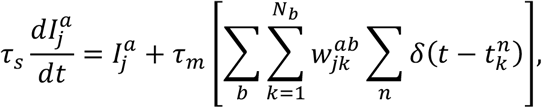

where *τ*_*s*_ is the synaptic time constant, 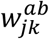 is the strength of the connection from neuron *k* in population *b* to neuron *j* in population *a* and 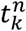 are the spike times of neuron *k*. The probability of a connection from population *b* to *a* is given by *p*_*a,b*,_ and if a connection exists, 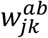 is set to either *w* or – *gw* for incoming excitatory or inhibitory inputs, respectively, and 0 otherwise.

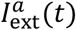 models the synaptic current from 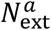 Poisson sources with connection strength *w* and firing rate 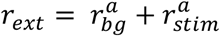, where 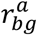 is the fixed background firing rate and 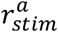 is the stimulus firing rate, which depends on the magnitude of the input stimulus (none, low, medium, or high). Instead of explicitly modeling the spiking behavior of this source, we make use of a diffusion approximation

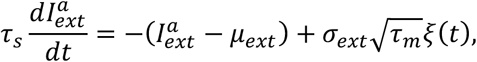

with

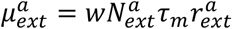, and

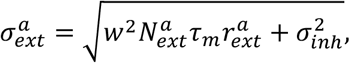

where 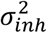 is a fixed level of background noise that accounts for additional variability from inhibitory inputs.

### Modeling noise-induced damage and recovery

To model the noise-induced damage seen experimentally, we decrease both the background and stimulus-related firing rates by factors *γ, β*^*a*^ < 1, so that the external firing rate becomes

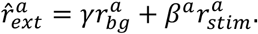

Recovery was modeled as a depolarizing or hyperpolarizing current that adjusted the resting potential directly,

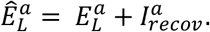

All parameter values for the LIF model can be found in Tables 1 and 2.

### Mean-field and diffusion approximation

Using the results from [3], we make a diffusion approximation for the recurrent inputs to a neuron. This approximation assumes that the input spike trains follow a Poisson distribution, are uncorrelated, and the amplitude of the depolarization due to each input it small 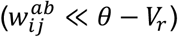.

Let *N*_*a*_ denote the size of population *a* and *P*_*a,b*_ denote the connection probability of a neuron in population *b* to a neuron in population *a*. The average number of incoming connections is therefore given by

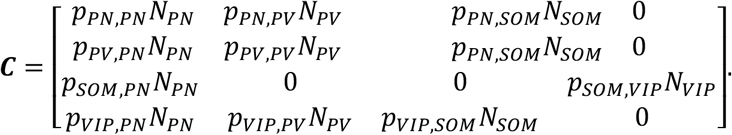

Letting *w* be a matrix of connection strengths

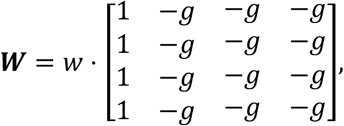

it follows that the average connectivity between populations is described by

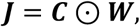

where ⊙ denotes the Hadamard product.

Denoting the steady-state firing rates as 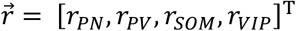, we define

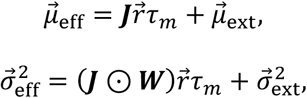

and the mean-field neuronal dynamics are described by the following system of stochastic differential equations

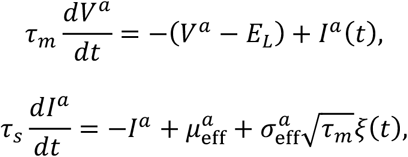

where *ξ* is a white noise term, with zero mean and unit variance density. It follows by [3] that up\ to the first order in 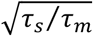, the steady state firing rates are given by

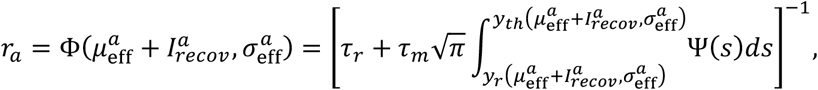

where

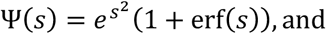

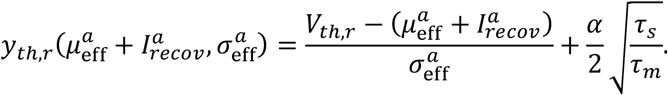

Here, 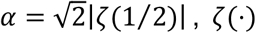 is the Riemann zeta function. Further, the population firing rate dynamics can be described by the following Wilson-Cowan equation

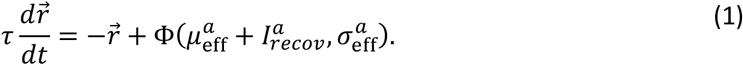

### Parameter sweep and stability criteria

For the three-population model, we estimated the firing rate of the excitatory population (*r*_*PN*_) at four stimulus levels (*r*_*stim*_) using the mean-field theory and took the excitatory gain to be the slope of the line of best fit 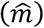

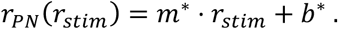

We then perform an extensive sweep in the 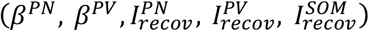 parameter space consisting of 225,000 possible parameter sets, estimating the firing rate and gain in the same way as the default case.

A parameter set 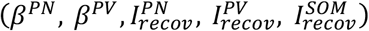 with gain 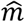 was deemed viable if it demonstrates the following traits observed in the experimental data:

1. 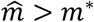
2. 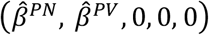, meaning the parameter set without recovery, show decrease gain from the default value *m*^∗^
3. The firing rate of all populations monotonically increased with stimulus strength
4. A low PN response threshold, defined to be >1Hz for the low stimulus value
5. Stable dynamics (see details below)

Some parameter sets were immediately discarded due to the stability criteria because the mean-field theory failed to converge to a stable solution. However, for some of the considered parameter sets, we found examples where the converged mean-field theory disagreed with the corresponding result from the spiking model, due to the spiking model being pushed into an unstable, synchronous, and heavily correlated regime. Due to this disagreement, we wanted to discard such parameter sets from the analysis.

As stated previously, the mean-field theory assumes that the spiking dynamics lays in an asynchronous regime, and this disagreement arises when this assumption breaks down. Unfortunately, it is not straight forward to use the theory to predict exactly when this disagreement will occur for a given parameter set^68,112^. Here, we use the eigenvalues of the Jacobian of the deterministic model to provide a conservative and unbiased threshold to disregard such parameter sets. Specifically, we linearize Eq (1) around the fixed point

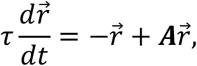

and then disregard parameter sets where max ℜ (*λ* (*A* − *I*)) = *k* > *k*_*thres*_. The deterministic system is unstable when *k* > 0, but in order to discard parameter sets that lead to the disagreement between the theory and the spiking model described above, we consider the conservative threshold of *k*_*thres*_ = −0.7.

The methods and criteria are the same for the four-population model, but having eliminated intrinsic changes in the SOM population experimentally and adding the VIP population to the model, we consider the parameter space 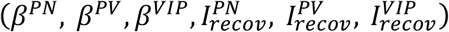. Following the results of the first model, the parameter space hypercube was also adjusted to only consider positive values of 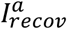. In total, 15,625 total parameter sets were considered.

### Numerical details

The spiking network is implemented Euler’s method with a timestep of 0.01 ms. Each trial consisted of 5 seconds, and the steady state firing rates were computed after averaging the spiking activity of the neurons across each population after discarding the first second of each trial. The firing rates for the mean-field theory were found via fixed point iteration of Eq (1), which halted when 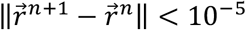.

### Statistics

For statistical comparisons between two independent groups that passed the Shapiro– Wilk normality test, we used unpaired t-tests. Otherwise, we used the Mann–Whitney rank-sum test, Kruskal–Wallis one-way analysis of variance, or Friedman test for non-normally distributed data. For comparisons between multiple groups having within-subject factors, a repeated measures two-way ANOVA test was used, Bonferroni corrections were used for multiple two-sample post hoc comparisons among sample groups; the significance level (α = 0.05) of the test was corrected via scaling by the reciprocal of the number of comparisons. Greenhouse-Geisser correction was used when the assumption of sphericity was violated. A permutation test (Wasserman, 2004) was used for two sample comparisons. Samples for which 5000 of 100,000 random permutations of the data resulted in mean differences greater than the observed difference in sample means were considered significant (p, 0.05). Significance levels are denoted as ∗, P <0.05. The details of statistical tests are described in the figure legends. Group data are presented as mean ± SEM. Sample sizes were not predetermined by statistical methods but were based on those commonly used in the field.

### Rigor and Transparency

Behavioral, *in vitro* electrophysiology, and histology experiments were conducted and analyzed in a blind mode regarding noise- and sham-exposed conditions. For *in vivo* imaging experiments, analysis was also done in a blind mode. Although the experimenter was not “blind” during the acquisition of the *in vivo* imaging experiments, those experiments involved identical and automated signal detection, inclusion, and analysis for both noise- and sham-exposed mice (as described in Methods). Thus, the experimenter did not have any influence over the experiment. Thus, between the data acquisition and analyses, all experiments and analyses are transparent, rigorous, and reproducible.

## Code and Data availability

Source data for all figures will be made available from the corresponding author upon reasonable request. The code is written in a combination of C and MATLAB, and those corresponding to the main results can be found on GitHub (https://github.com/gregoryhandy) upon acceptance for publication. Other custom Matlab codes used in this study will be made available from the corresponding author upon reasonable request.

## Acknowledgments

We thank Dr. Patrick Cody for help in data analysis.

## Author contributions

M.K, and T.T. designed the study. M.K performed *in vivo* experiments and analyzed data. G.H. and B.D. designed and programmed the computational modelling. S.K. performed electrophysiology experiments. L.L.B performed behavioral experiments. B.B performed cochlear histology. M.K., G.H., B.D., and T.T. wrote the manuscript.

## Figure Legends

**Figure 1 supplement.**
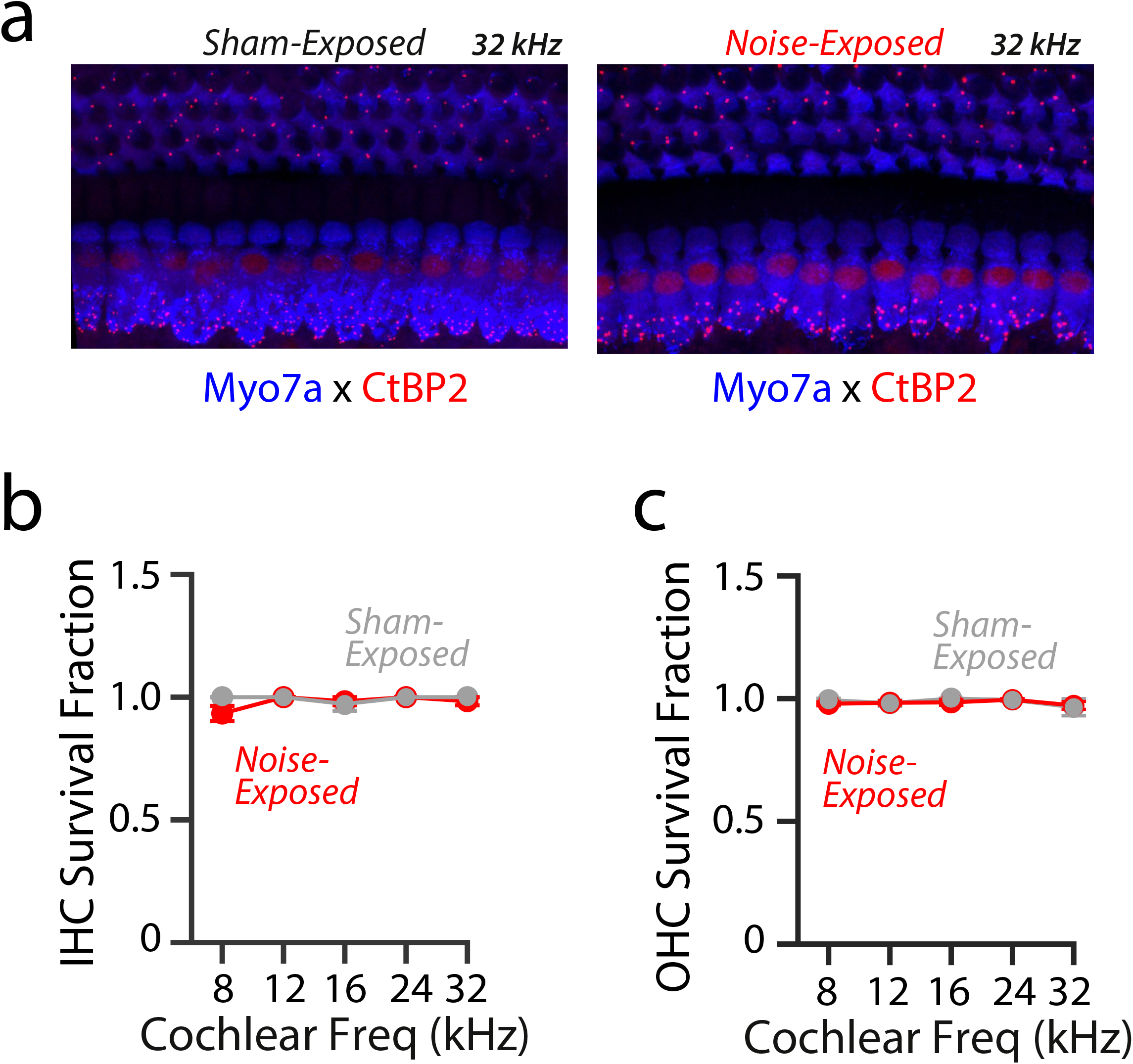
Noise exposure did not alter the IHC and OHC structures. **a)** Representative images of OHCs from the 32 kHz region of sham- (left) and noise- (right) exposed mice. **b)** Quantification of IHC survival from sham- (grey) and noise- (red) exposed mice. (Noise: 5 mice vs. sham: 4 mice, 2-way ANOVA; exposure x frequency, F = 1.89, p = 0.13; effect of exposure, F = 2.7, p = 0.10). **c)** Quantification of OHC survival from sham- (grey) and noise- (red) exposed mice. (Noise: 5 mice vs. sham: 4 mice, 2-way ANOVA; exposure x frequency, F = 0.26, p = 0.89; effect of exposure, F = 0.81, p = 0.37).

**Figure 2 supplement.**
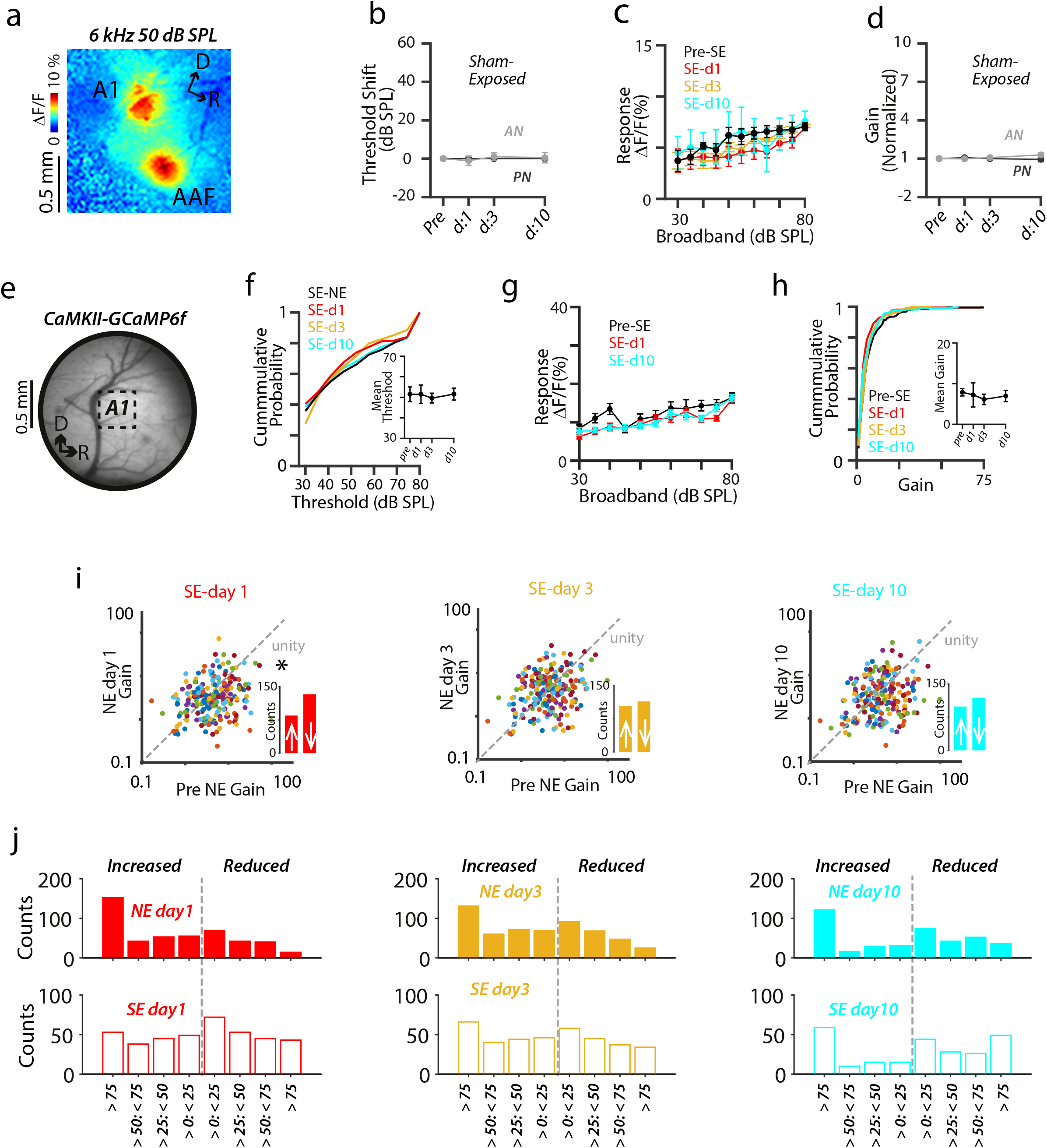
Sham exposure did not alter the sound-evoked activity of A1 L2/3 PN neurons. **(a)** Localization of A1. A 6 kHz 50 dB SPL tone triggered GCaMP6s fluorescence responses in two regions of the auditory cortex representing A1 and the anterior auditory field (AAF; D stands dorsal and R for rostral). **(b)** Average change in response thresholds of A1 PNs (dark grey) at 1, 3, and 10 days after sham exposure. (n = 3 mice, 1-way repeated measure ANOVA, F = 0, p > 0.99). Average change in AN threshold (light grey) reproduced from **Figure 1. (c)** Average sound-evoked responses of A1 PNs to broadband sounds from sham-exposed mice. (n = 3 mice, 2-way ANOVA; time x sound level interaction, F = 1.3, p = 0.35; effect of time, F = 0.54, p = 0.56). **(d)** Average response gain of A1 PNs (dark grey) normalized to pre-sham-exposed gain after sham exposure at 1, 3, and 10 days. (n = 3 mice, 1-way repeated measure ANOVA, F = 1.3, p = 0.34). Normalized AN gain (light grey) reproduced from **Figure 1. e)**. Implantation of cranial glass window over A1. **(f)** Cumulative response threshold of A1 L2/3 PNs, before and after sham exposure. Inset: Average mean threshold of PNs per mouse (n = 5 mice, 1-way repeated measure ANOVA, F = 0.17, p = 0.87). **(g)** Average sound-evoked responses of A1 L2/3 individual PNs to broadband sounds from sham-exposed mice. (2-way ANOVA; sound intensity and time interaction, F = 1.6, p = 0.072; effect of time, F = 2.9, p = 0.065). **(h)** Cumulative gain of A1 L2/3 PNs, before and after sham exposure. Inset: Average mean gain of PNs per mouse (1-way repeated measure ANOVA, F = 0.29, p = 0.71). **(p)** Scatter plots of the gain of individual A1 L2/3 PNs before and after sham exposure. Dotted line represents unity. Insets: Bar graphs representing the number of neurons showing increased gain (↑ above unity) and reduced gain (↓ below unity) after NIHL. PreSE vs. SEday1: p = 0.006, PreSE vs. SEday3: p = 0.11, and PreSE vs. SEday10: p = 0.06; permutation test. **j)** Histograms showing percentage changes in the gain of L2/3 PNs after noise (top) and sham (bottom) exposure.

**Figure 4 supplement.**
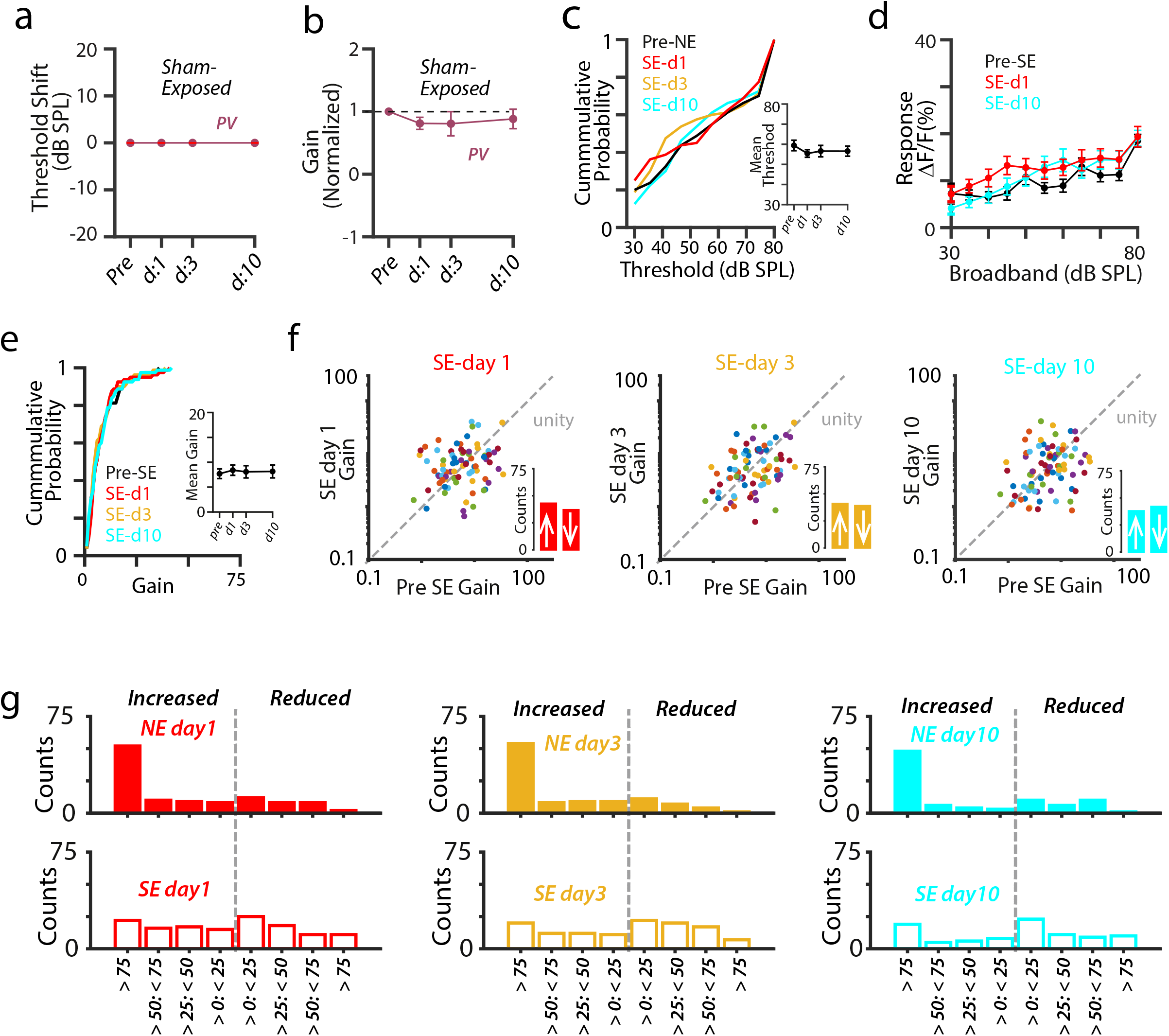
Sham exposure did not alter the sound-evoked activity of A1 L2/3 PV neurons. **(a)** Average change in response thresholds of A1 PV neurons at 1, 3, and 10 days after sham exposure. (n = 3 mice, 1-way repeated measure ANOVA, F = 0, p > 0.99). **(b)** Average response gain of A1 PV neurons normalized to pre-sham-exposed gain after sham exposure at 1, 3, and 10 days. (n = 3 mice, 1-way repeated measure ANOVA, F = 1.09, p = 0.40). **(c)** Cumulative response threshold of A1 L2/3 PV neurons, before and after sham-exposure. Inset: Average mean threshold of PV neurons per mouse (n = 8 mice, 1-way repeated measure ANOVA, F = 0.79, p = 0.46). **(d)** Average sound-evoked responses of A1 L2/3 individual PV neurons to broadband sounds from sham-exposed mice. (2-way ANOVA; sound intensity and time interaction, F = 1.3, p = 0.16; effect of time, F = 1.6, p = 0.16). **(e)** Cumulative gain of A1 L2/3 PV neurons, before and after sham exposure. Inset: Average mean gain of PV neurons per mouse (1-way repeated measure ANOVA, F = 0.14, p = 0.86). **(f)** Scatter plots of the gain of individual A1 L2/3 PV neurons, before and after sham exposure. Dotted line represents unity. Insets: Bar graphs representing number of neurons showing increased gain (↑ above unity) and reduced gain (↓ below unity) after NIHL. PreSE vs. SEday1: p = 0.98, PreSE vs. SEday3: p = 0.67, and PreSE vs. SEday10: p = 0.96; permutation test. **g)** Histograms showing percentage changes in the gain of L2/3 PV neurons after noise (top) and sham (bottom) exposure.

**Figure 5 supplement.**
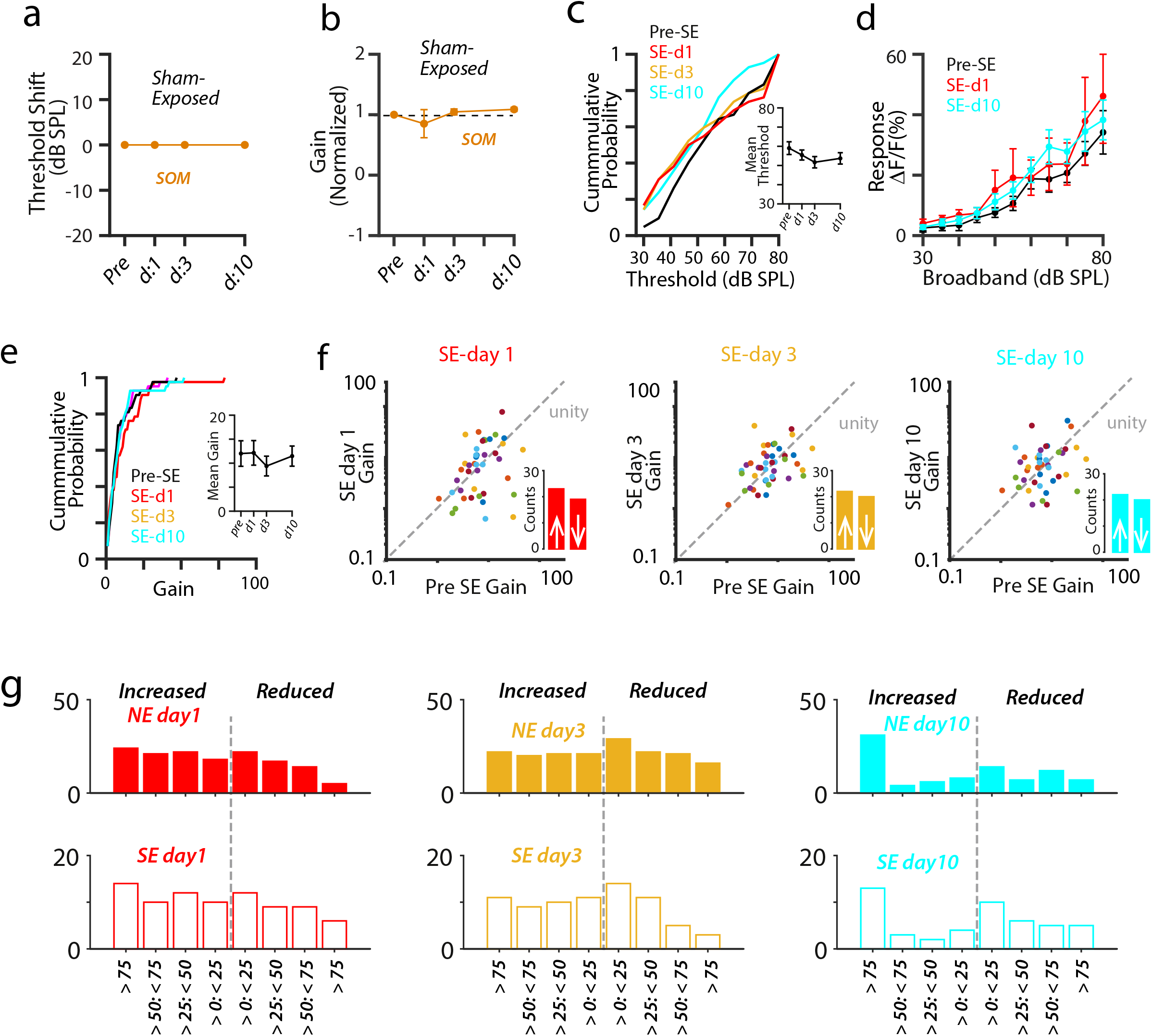
Sham exposure did not alter the sound-evoked activity of A1 L2/3 SOM neurons. **(a)** Average change in response thresholds of A1 SOM neurons at 1, 3, and 10 days after sham exposure. (n = 4 mice, 1-way repeated measure ANOVA, F = 0, p > 0.99). **(b)** Average response gain of A1 SOM neurons normalized to pre-sham-exposed gain after sham exposure at 1, 3, and 10 days. (n = 4 mice, 1-way repeated measure ANOVA, F = 0.85, p = 0.42). **(c)** Cumulative response threshold of A1 L2/3 SOM neurons, before and after sham exposure. Inset: Average mean threshold of SOM neurons per mouse (n = 9 mice, 1-way repeated measure ANOVA, F = 1.1, p = 0.33). **(d)** Average sound-evoked responses of A1 L2/3 individual SOM neurons to broadband sounds from sham-exposed mice. (2-way ANOVA; sound intensity and time interaction, F = 0.92, p = 0.43; effect of time, F = 0.62, p = 0.49). **(e)** Cumulative gain of A1 L2/3 SOM neurons, before and after sham exposure. Inset: Average mean gain of SOM neurons per mouse (1-way repeated measure ANOVA, F = 0.35, p = 0.70). **(f)** Scatter plots of the gain of individual A1 L2/3 SOM neurons, before and after sham exposure. Dotted line represents unity. Insets: Bar graphs representing the number of neurons showing increased gain (↑ above unity) and reduced gain (↓ below unity) after NIHL. PreSE vs. SEday1: p = 0.55, PreSE vs. SEday3: p = 0.94, and PreSE vs. SEday10: p = 0.89; permutation test. **g)** Histograms showing percentage changes in the gain of L2/3 SOM neurons after noise (top) and sham (bottom) exposure.

**Figure 8 supplement.**
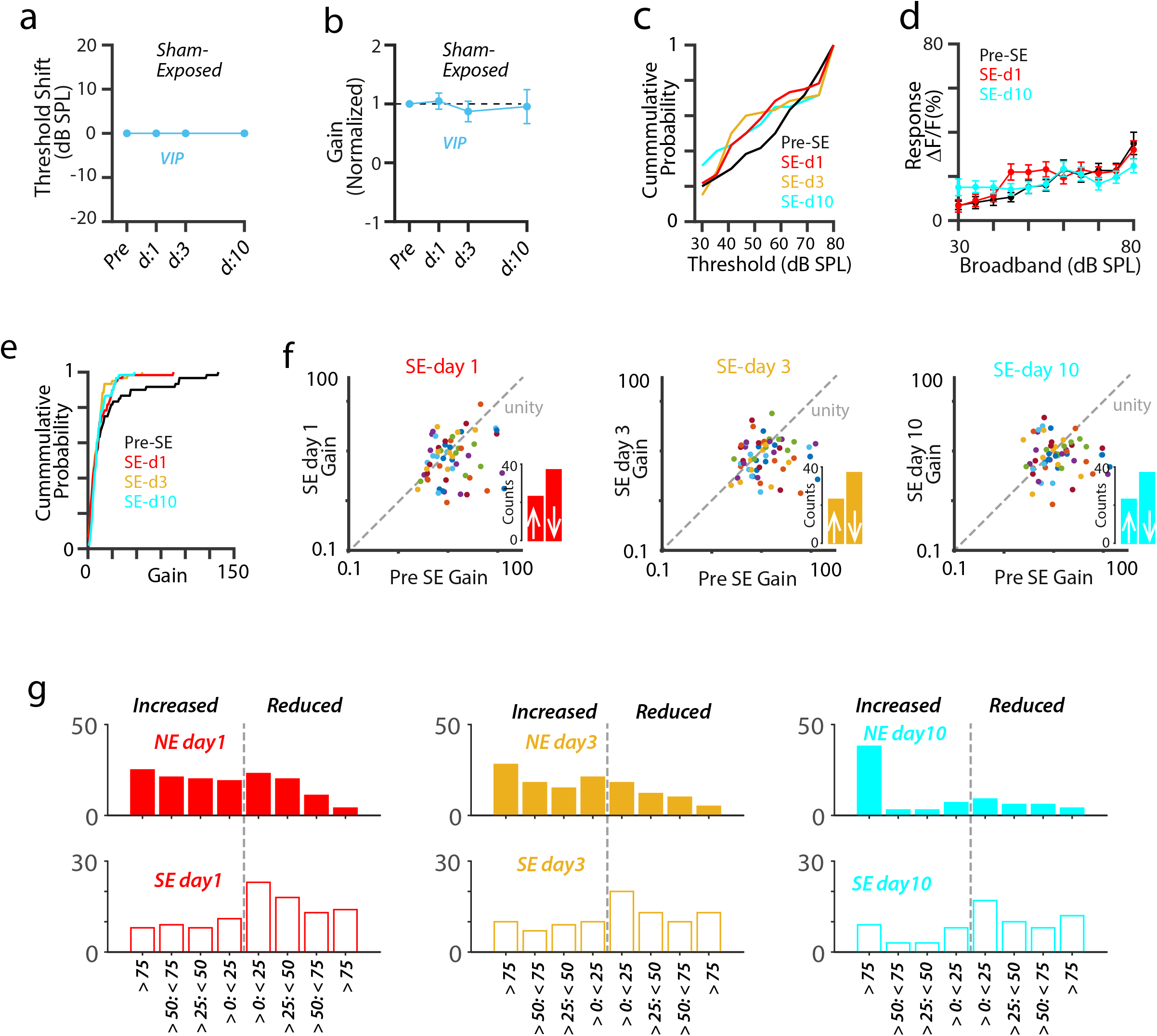
Sham exposure reduced the gain of A1 L2/3 VIP neurons. **(a)** Average change in response thresholds of A1 VIP neurons at 1, 3, and 10 days after sham exposure. (n = 3 mice, 1-way repeated measure ANOVA, F = 0, p > 0.99). **(b)** Average response gain of A1 VIP neurons normalized to pre-sham-exposed gain after sham exposure at 1, 3, and 10 days. (n = 3 mice, 1-way repeated measure ANOVA, F = 0.03, p = 0.98). **(c)** Cumulative response threshold of A1 L2/3 VIP neurons, before and after sham-exposure. Inset: Average mean threshold of VIP neurons per mouse (n = 6 mice, 1-way repeated measure ANOVA, F = 0.52, p = 0.62). **(d)** Average sound-evoked responses of A1 L2/3 individual VIP neurons to broadband sounds from sham-exposed mice. (2-way ANOVA; sound intensity and time interaction, F = 4.5, p = 0.01; effect of time, F = 0.49, p = 0.69). **(e)** Cumulative gain of A1 L2/3 VIP neurons, before and after sham exposure. Inset: Average mean gain of VIP neurons per mouse (1-way repeated measure ANOVA, F = 0.44, p = 0.61). **(f)** Scatter plots of the gain of individual A1 L2/3 VIP neurons, before and after sham exposure. Dotted line represents unity. Insets: Bar graphs representing number of neurons showing increased gain (↑ above unity) and reduced gain (↓ below unity) after NIHL. PreSE vs. SEday1: p = 0.01, PreSE vs. SEday3: p = 0.005, and PreSE vs. SEday10: p = 0.006; permutation test. **g)** Histograms showing percentage changes in the gain of L2/3 VIP neurons after noise (top) and sham (bottom) exposure.

